# Integrating gene expression, spatial location and histology to identify spatial domains and spatially variable genes by graph convolutional network

**DOI:** 10.1101/2020.11.30.405118

**Authors:** Jian Hu, Xiangjie Li, Kyle Coleman, Amelia Schroeder, David J. Irwin, Edward B. Lee, Russell T. Shinohara, Mingyao Li

**Author notes:** **Correspondence:** Jian Hu, Mingyao Li.

## Abstract

Recent advances in spatial transcriptomics technologies have enabled comprehensive characterization of gene expression patterns in the context of tissue microenvironment. To elucidate spatial gene expression variation, we present SpaGCN, a graph convolutional network approach that integrates gene expression, spatial location and histology in spatial transcriptomics data analysis. Through graph convolution, SpaGCN aggregates gene expression of each spot from its neighboring spots, which enables the identification of spatial domains with coherent expression and histology. The subsequent domain guided differential expression analysis then detects genes with enriched expression patterns in the identified domains. Analyzing five spatially resolved transcriptomics datasets using SpaGCN, we show it can detect genes with much more enriched spatial expression patterns than existing methods. Furthermore, genes detected by SpaGCN are transferrable and can be utilized to study spatial variation of gene expression in other datasets. SpaGCN is computationally fast, making it a desirable tool for spatial transcriptomics studies.

## Introduction

Recent advances in spatial transcriptomics technologies have enabled gene expression profiling with spatial information in tissues^1^. Knowledge of the relative locations of different cells in a tissue is critical for understanding disease pathology because spatial information helps in understanding how the gene expression of a cell is influenced by its surrounding environment and how neighboring regions interact at the gene expression level. Experimental methods to generate spatial transcriptomics data can be broadly classified into two categories: 1) single-molecule fluorescence *in situ* hybridization (smFISH) based techniques, such as MERFISH^2^ and seqFISH^3^, which measure expression level for hundreds of genes with subcellular spatial resolution in a single cell; and 2) spatial barcoding followed by next generation sequencing based techniques, such as SLIDE-seq^4^ and 10X Genomics Visium, which measure the expression level for thousands of genes in captured locations, referred to as spots. These different spatial transcriptomics techniques have made it possible to uncover the complex transcriptional architecture of heterogenous tissues and enhanced our understanding of cellular mechanisms in diseases^5,6^.

In spatial transcriptomics studies, an important step is identifying spatial domains defined as regions that are spatially coherent in both gene expression and histology. Identifying spatial domains requires methods that can jointly consider gene expression, spatial location, and histology. Traditional clustering methods such as K-means and Louvain’s method^7^ can only take gene expression data as input, and the resulting clusters may not be contiguous due to the lack of consideration of spatial information and histology. To account for spatial dependency of gene expression, new methods have been developed. For example, stLearn^8^ uses features extracted from histology image as well as expression of neighboring spots to spatially smooth gene expression data before clustering; BayesSpace^9^ employs a Bayesian approach for clustering analysis by imposing a prior that gives higher weight to spots that are physically close; Zhu *et al*.^10^ uses a Hidden-Markov random field approach to model spatial dependency of gene expression. Although these methods can cluster spots or cells into distinct groups, they do not provide biological interpretations of the identified spatial domains.

To link spatial domains with biological functions at the gene expression level, it is crucial to identify genes that show enriched expression in the identified domains. Due to spatial variation of cell types in tissue, the difference of gene expression between different domains is mainly driven by cell type composition variation. On the other hand, information on spatial location and the corresponding histology allows the construction of an anatomy-based taxonomy of the tissue, which provides a useful perspective on cell type composition. Although stLearns integrates gene expression, spatial location, and histology information in clustering, the putative correspondence between cell type difference and organizational structure of the tissue remains unclear. As reported in Saiselet *et al*.^11^, many spatial regions are highly intermixed in terms of cell types. Without further downstream gene-level analysis, the spatial domains detected by stLearn still suffer from the lack of interpretability. Recently, new methods such as Trendsceek^12^, SpatialDE^13^, and SPARK^14^ have been developed to detect spatially variable genes (SVGs). These methods examine each gene independently and return a p-value to represent the spatial variability of a gene. However, due to the lack of consideration of tissue taxonomy, genes detected by these methods do not have a guaranteed spatial expression pattern, making it difficult to utilize these genes for further biological investigations.

Rather than considering spatial domain identification and SVG detection as separate problems, we developed SpaGCN, a graph convolutional network-based approach that considers these two problems jointly. Using a graph convolutional network with an added iterative clustering layer, SpaGCN first identifies spatial domains by integrating gene expression, spatial location, and histology together through the construction of an undirected weighted graph that represents the spatial dependency of the data. For each spatial domain, SpaGCN then detects SVGs that are enriched in the domain against its surrounding regions by differential expression analysis guided by domain information. SpaGCN also has the option to detect meta genes that are uniquely expressed in a given domain. The spatial domains and the corresponding SVGs and meta genes detected for these domains provide a comprehensive picture on the spatial gradients in gene expression in tissue.

## Results

### Overview of SpaGCN and evaluation

SpaGCN is applicable to both sequencing-based and smFISH-based data. As shown in Fig. 1a, SpaGCN first builds a graph to represent the relationship of all samples (spots in sequencing-based or cells in smFISH-based data) considering both spatial location and histology information. Next, SpaGCN utilizes a graph convolutional layer to aggregate gene expression information from neighboring samples. Then, SpaGCN uses the aggregated gene expression matrix to cluster samples using an unsupervised iterative clustering algorithm^15^. Each cluster is considered as a spatial domain from which SpaGCN then detects SVGs that are enriched in a domain by differential expression analysis (Fig. 1b). When a single gene cannot mark expression pattern of a spatial domain, SpaGCN will construct a meta gene, formed by the combination of multiple SVGs, to represent gene expression of the domain. Since the expression profile of a spot/cell is heavily influenced by its local microenvironment, SpaGCN also offers the option of subcluster detection within each spatial domain. SVGs can also be detected to help in understanding the function of each sub-spatial domain.

**Figure 1.**
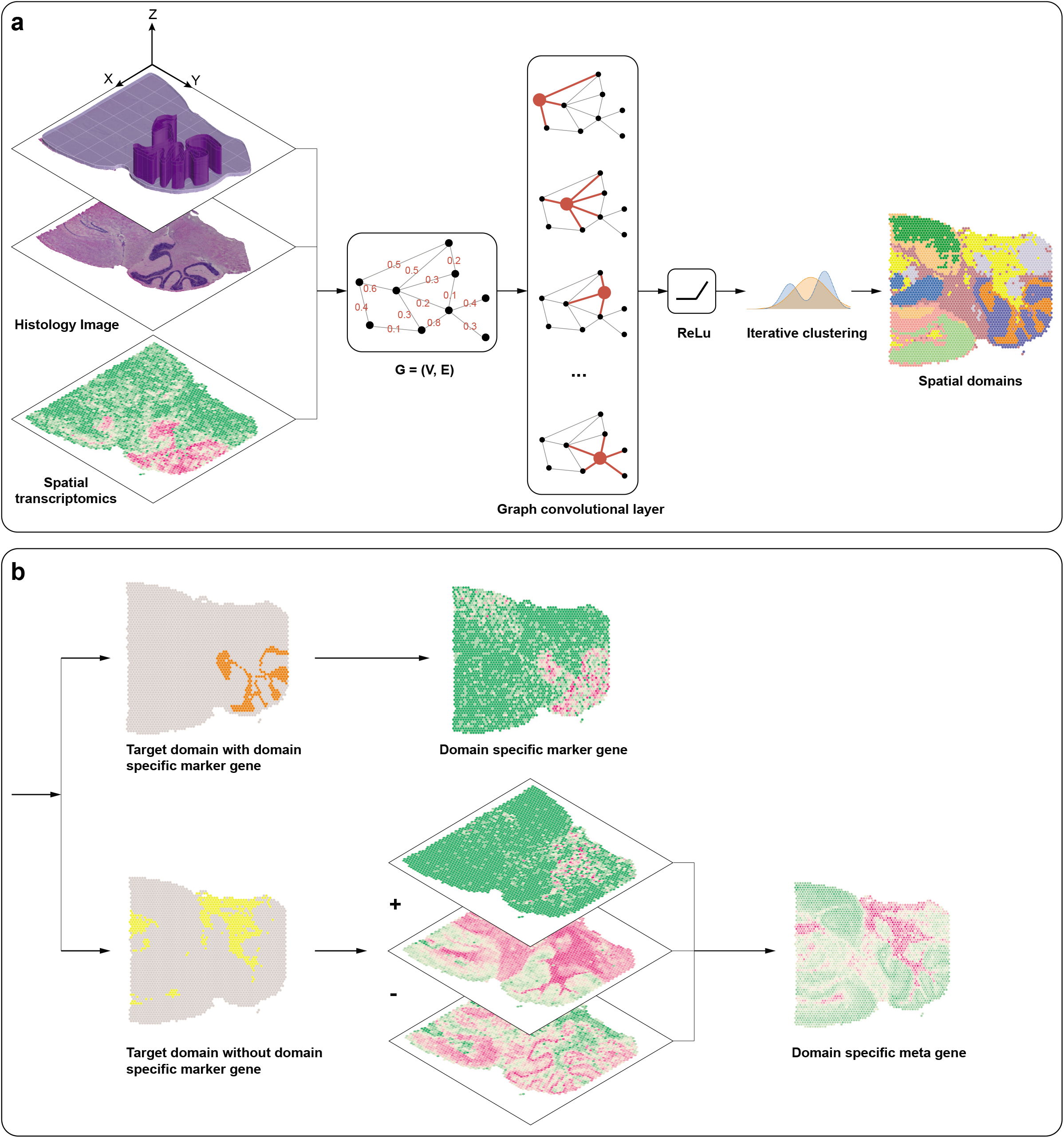
Workflow of SpaGCN. **a**, SpaGCN starts from integrating gene expression, spatial location and histology information using a graph convolutional network (GCN), then separates spots into different spatial domains using unsupervised iterative clustering. The GCN is based on an undirected weighted graph in which the edge weight between every two spots is determined by Euclidean distance between the two spots, defined by the spatial coordinates (*x, y*) and the 3-rd dimensional coordinate *z*, obtained from the RGB values in the histology image. **b**, For each detected spatial domain, SpaGCN identifies SVGs or meta genes by domain guided differential expression analysis.

To showcase the strength and scalability of SpaGCN, we applied it to five publicly available datasets, including four datasets generated by sequencing-based techniques and one dataset generated by MERFISH (Supplementary Table 1). The spatial domains identified by SpaGCN agree better with known tissue layer structure than K-means and Louvain’s clustering. We also compared SVGs detected by SpaGCN with those detected by SPARK^14^ and SpatialDE^13^, and found that the SVGs detected by SpaGCN have more coherent expression patterns and better biological interpretability than the other two methods. The specificity of spatial expression patterns revealed by SpaGCN detected SVGs were further confirmed by Moran’s *I* statistic^16^, a metric that quantifies the spatial autocorrelation of detected genes.

### Application to mouse olfactory bulb data

To evaluate the performance of SpaGCN, we first analyzed a mouse olfactory bulb (MOB) dataset^17^, which consists of 16,218 genes measured in 262 spots. The main olfactory bulb has five layers, ordered from surface to the center as follows: glomerular layer, external plexiform layer, mitral cell layer, internal plexiform layer, and granule cell layer. We compared SpaGCN’s clustering results to K-means and Louvain by setting the number of clusters at 5 for all three methods. As shown in Fig. 2a, K-means only identified 3 main spatial domains, with only few spots assigned to domains 1 and 3. Louvain’s method identified 5 main spatial domains. However, since it does not consider spatial and histology information, domains 2, 3, and 4 have blurred boundaries and more outliers than SpaGCN. By contrast, the domains detected by SpaGCN agree better with the biologically known 5-layer structure of the MOB.

**Figure 2.**
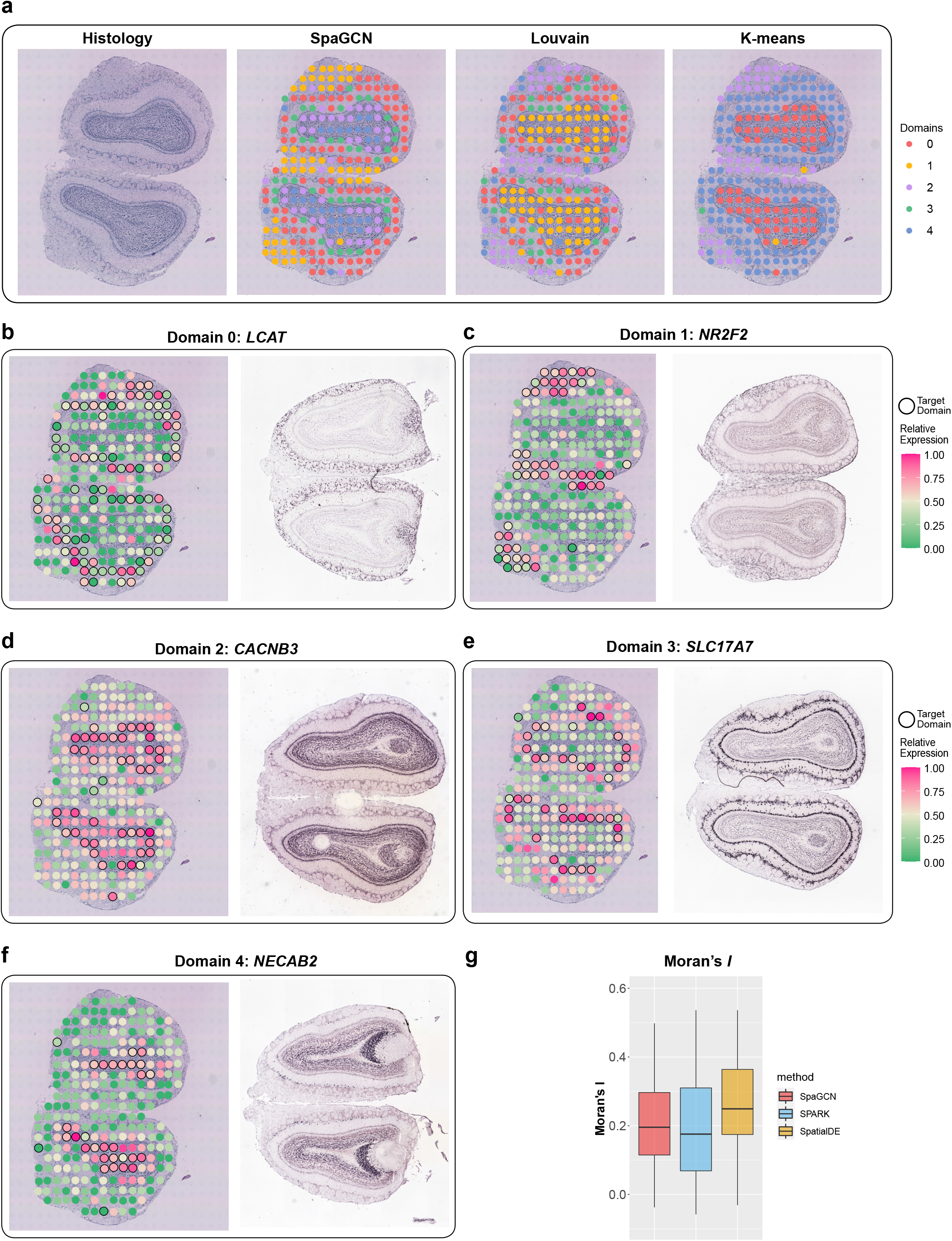
Spatial domains and SVGs detected in the mouse olfactory bulb dataset. **a**, Histology image of the tissue section and spatial domains detected by SpaGCN, Louvain’s method, and K-means clustering. **b-f**, Spatial expression patterns of SVGs detected by SpaGCN for domains 0 (*LCAT*), 1 (*NR2F2*), 2 (*CACNB3*),3 (*SLC17A7*), and 4 (*NECAB2*), and the corresponding *in situ* hybridization of these SVGs obtained from the Allen Brain Atlas. **g**, Boxplot of Moran’s *I* values for SVGs detected by SpaGCN, SPARK, and SpatialDE.

To understand the functions of the SpaGCN identified spatial domains, we next detected SVGs for each spatial domain. In total, SpaGCN detected 60 SVGs. Fig. 2b-f shows a randomly selected SVG for each domain, and all genes show strong specificity for the corresponding domain. The *In Situ* Hybridization labelling of these genes from the Allen Brain Institute further confirmed the correspondence of the spatial domains detected by SpaGCN. Additional SVGs detected by SpaGCN are shown in Supplementary Fig. 1. As a comparison, we also detected SVGs using SpatialDE and SPARK. SpatialDE identified 67 SVGs, but only 12 of them overlapped with SpaGCN results (Supplementary Fig. 2). We further looked into the 55 genes detected exclusively by SpatialDE and found many of the genes are expressed in only a few spots or are highly expressed in most of the spots, leading to false detections of significant spatial patterns (Supplementary Fig. 3). By contrast, SpaGCN avoided this issue by filtering out genes using minimum within group expression fraction and maximum between group expression fraction. SPARK detected 772 genes, with 49 overlapping with SapGCN (Supplementary Fig. 2). However, we found that the SPARK results indicate that 274 genes have FDR-adjusted p-values less than 0.00001 with 14 of them having the smallest identical FDR-adjusted p-value of 4.42e-13. As a result, the SPARK p-values are not informative in differentiating the degree of spatial variability between different genes. Of note, none of these 14 genes were detected by SpaGCN. Further examination revealed that some of these genes show spatial variability, but more than half of them are only expressed in a few spots or highly expressed in most of the spots (Supplementary Fig. 4). The FDR-adjusted p-value distribution of SPARK and q-value distribution of SpatialDE are highly skewed toward 0, making it challenging to select informative SVGs based on their p-values or q-values alone (Supplementary Fig. 5).

To compare SVGs detected by different methods quantitatively, we calculated the Moran’s *I* statistic, which measures the spatial autocorrelation for each gene. Fig. 2g shows the distribution of Moran’s *I*. Although all SpaGCN detected SVGs have clear spatial patterns, their Moran’s *I* values are not significantly higher than the SVGs detected by SPARK and SpatialDE (median of 0.20 for SpaGCN against 0.18 for SPARK and 0.25 for SpatialDE). Further examination revealed that many SVGs detected by SPARK and SpatialDE are expressed in multiple adjacent spatial domains. For example, the gene *PCP4* uniquely detected by SpatialDE is expressed in two adjacent layers (domains 2 and 4 defined by SpaGCN) (Supplementary Fig. 6). By contrast, all the SVGs detected by SpaGCN are domain specific, offering interpretation in alignment with our knowledge of layer structure. We note that less informative SVGs with clear, but non-domain specific, spatial patterns, such as *PCP4*, can also be detected by SpaGCN if the user combines domains 2 and 4 as the target domain in SVG detection.

### Application to mouse posterior brain data

Next, we analyzed a dataset generated from mouse posterior cerebrum, cerebellum and brainstem by 10X Genomics that includes 3,353 spots and 31,053 genes^18^. We compared the clustering results of SpaGCN with K-means and Louvain’s clustering. The number of clusters in K-means and resolution in Louvain were set to generate the same number of clusters as SpaGCN (10 clusters). Fig. 3a shows that Louvain’s clustering is similar to SpaGCN, but the spatial domains detected by SpaGCN are more spatially contiguous than Louvain’s results. The integrity of SpaGCN’s spatial domains stems from the aggregation of gene expression based on spatial information and histology, which ensures that the genes detected by differential expression analysis have clear spatial expression patterns.

**Figure 3.**
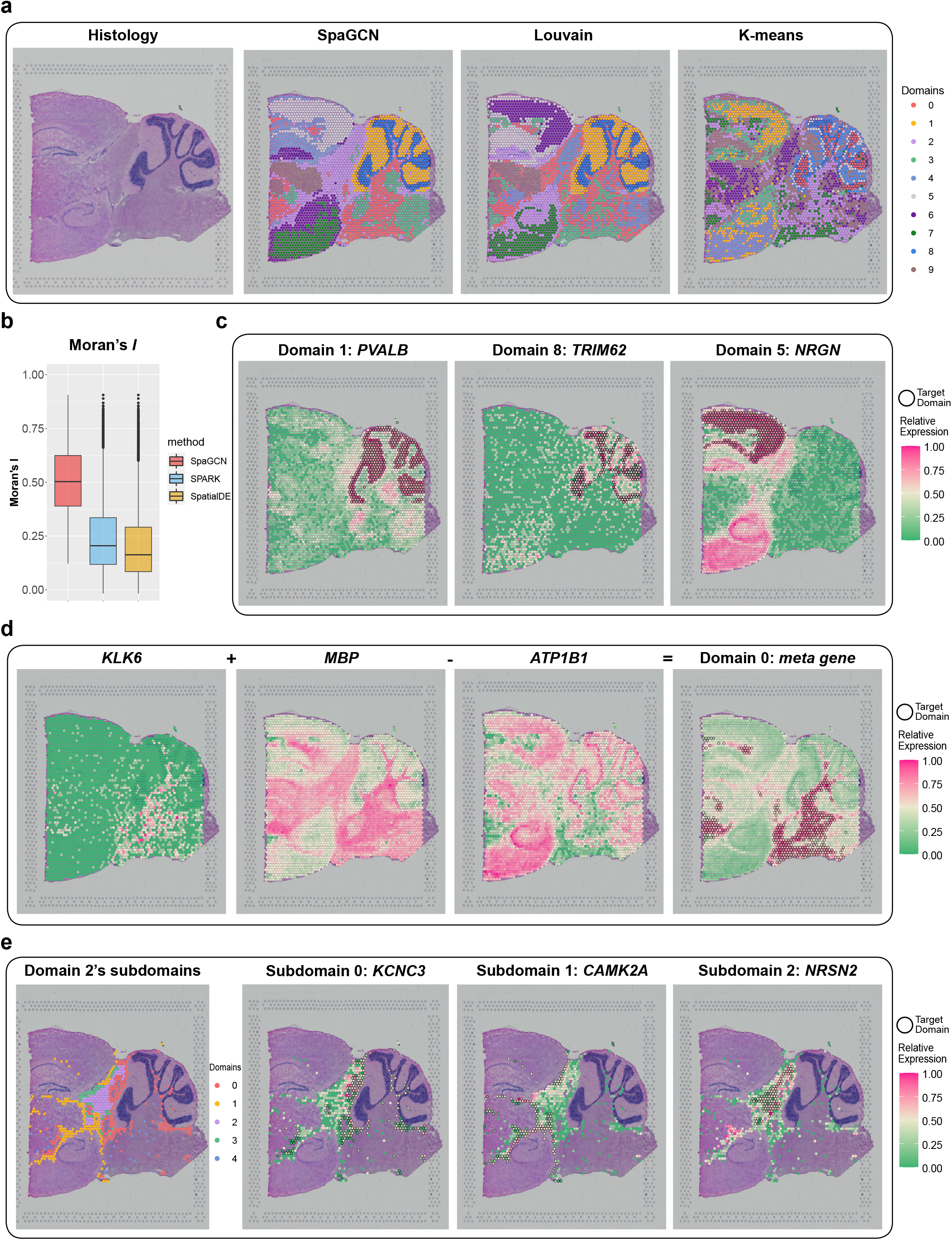
Spatial domains and SVGs detected in the mouse brain posterior brain dataset. **a**, Histology image of the tissue section and spatial domains detected by SpaGCN, Louvain’s method, and K-means clustering. **b**, Boxplot of Moran’s *I* values for SVGs detected by SpaGCN, SPARK, and SpatialDE. **c**, Spatial expression patterns of SVGs detected by SpaGCN for domain 1 (*PVALB*), 8 (*TRIM62*), and 5 (*NRGN*). **d**, Spatial expression patterns of genes *KLK6, MBP, ATP1B1*, which form the specific marker meta gene for domain 0 (*KLK6 + MBP - ATP1B1*). **e**, Clustering results for 5 sub-domains detected by SpaGCN for domain 2, and the spatial expression patterns of SVGs for sub-domains 0 (*KCNC3*), 1 (*CAMK2A*), and 2 (*NRSN2*).

SpaGCN detected 523 SVGs for the 10 spatial domains while SPARK and SpatialDE detected 9,678 and 12,676 SVGs, respectively (Supplementary Fig. 7). We hypothesized that the substantially larger number of SVGs detected by SPARK and SpatialDE are due to the lack of spatial expression patterns that exist in the data. To confirm this hypothesis, we calculated the Moran’s *I* statistic for all detected SVGs (Fig. 3b). The Moran’s *I* values of SpaGCN detected SVGs are much higher than those detected by SPARK and SpatialDE (median of 0.50 for SpaGCN against 0.21 for SPARK and 0.16 for SpatialDE). Closer examination of the SVGs detected by SPARK and SpatialDE revealed that most of the SVGs suffer from one of the two problems observed previously in the MOB dataset: they are (1) only expressed in a few spots or highly expressed in most of the spots, suggesting high false positive rates for SPARK and SpatialDE or (2) spatially variable, but expressed in multiple adjacent spatial domains, making it difficult to interpret. Another limitation of these two methods is that the FDR-adjusted p-value from SPARK and q-value from SpatialDE are not informative. Genes with similar p-values/q-values do not necessarily show similar spatial pattern and a smaller p-value/q-value does not guarantee a better spatial pattern (Supplementary Fig. 8 and Supplementary Fig. 9). The p-value and q-value distributions of SPARK and SpatialDE are highly skewed toward 0 (Supplementary Fig. 10). By contrast, the SVGs detected by SpaGCN were enriched in specific spatial domains (Supplementary Fig. 11) and their expression patterns are transferable to an adjacent tissue slice in the mouse posterior brain (Supplementary Fig. 12). Further, multiple domain adaptive filtering criteria implemented in SpaGCN allow it to eliminate false positive SVGs and ensure all detected SVGs have clear spatial expression patterns.

To illustrate why appropriate filtering is important, we use domains 1, 5, and 8 as an example. For each of these domains, SpaGCN detected a single SVG enriched in that region. As shown in Fig. 3c, *PVALB* is enriched in domain 1, and *TRM62* is enriched in domain 8. Although domains 1 and 8 are adjacent to each other, these two SVGs can still well mark these domains. *NRGN* is a SVG that SpaGCN detected for domains 5 and 7. The high expression of *NRGN* in domains 5 and 7 also indicate that these two domains are neuroanatomically similar – both consisting of cortex and the pyramidal layer of the hippocampus. Both the cortex and hippocampus are regions that are on the curved surface of the brain. This posterior brain tissue section has the top part of the curved surface in domain 5 and the bottom part of the curved surface in domain 7. Domains 5 and 7, which would be contiguous in a complete 3D reconstruction, are artifactually separated due to the way the section was cut. Therefore, it is not surprising that in addition to *NRGN*, SpaGCN also detected many other SVGs, such as *APP, ATP6V1G2, CALM2, CHN1, CLSTN1, ARPP21, CYP46A1, DCLK1, LINGO1*, and *MARCKS*, that are highly expressed in both domains 5 and 7 (Supplementary Fig. 11). The unique and powerful SVG detection procedure in SpaGCN ensures that genes like these are not missed.

SpaGCN did not identify any SVGs for domain 0. However, we reason that a meta gene, formed by the combination of multiple genes, may better reveal spatial patterns than any single genes. We used domain 0 as an example to show how SpaGCN can create informative meta genes to mark a spatial domain (Fig. 3d). First, by lowering the filtering thresholds, SpaGCN identified *KLK6* which is highly expressed in the lower part of domain 0. Using *KLK6* as a starting gene, SpaGCN used a novel approach to find a log-linear combination of gene expression of *KLK6, MBP* and *ATP1B1*, which accurately marked the spatial domain 0. In this meta gene, *KLK6* and *MBP* are considered as positive markers because they are highly expressed in some spots in domain 0, whereas *ATP1B1* is considered a negative marker as it is mainly expressed in regions other than domain 0. Previous studies have shown that *KLK6* and *MBP* expression is restricted to oligodendrocytes, while *ATP1B1* is mainly expressed in neurons and astrocytes^19^. This resonates the fact that domain 0 represents white matter which is dominated by oligodendrocytes and has few neuronal cell bodies. Therefore, the genes that make up this meta gene have meaningful biological interpretation. Using this meta gene detection procedure, we also detected meta genes for domains 2, 7, 8 and 9, and found that these meta genes are transferrable to an adjacent tissue slice (Supplementary Fig. 13).

The expression profile and biological function of a spot is heavily influenced by its neighboring spots. The surrounding spots can trigger a response pathway or signal the spot to perform certain tasks. Although the spots in one spatial domain detected by SpaGCN are spatially coherent and have similar gene expression patterns, they may still have different functions since their surrounding spots are different. For instance, spots located near the boundary of a spatial domain may have different functions compared to spots located in the inner part of the domain. To learn more about the effect of different neighborhoods on the spots, we performed sub-domain detection. For example, domain 2 is located in the center of the tissue slice and surrounded by multiple other spatial domains. As a result, the neighboring environment for spots in domain 2 varies. As shown in Fig. 3e, domain 2 was separated into 5 sub-domains which are located either in the center or different boundary regions of domain 2, suggesting that differences in the neighborhoods of spots contribute to within-domain heterogeneity. SVGs detected for each sub-domain can help us understand the gene expression variability of spots within each sub-domain.

### Application to LIBD human dorsolateral prefrontal cortex data

In addition to the datasets described previously, SpaGCN also showed advantage over competing methods when evaluated on the LIBD human dorsolateral prefrontal cortex (DLPFC) data^20^. The LIBD study sequenced 12 slices from DLPFC that spans six neuronal layers plus white matter. We started from analyzing slice 151673, which includes 3,639 spots and 33,538 genes. As the original publication manually annotated the tissue into 7 layers, for fair comparison, the number of clusters was also set at 7 for SpaGCN, K-means, and Louvain. As shown in Fig. 4a, K-means and Louvain failed to separate the tissue into layers with clear boundary. By contrast, SpaGCN successfully identified layer structures with clear boundaries. The Adjusted Rand Indexes (ARIs) for the SpaGCN, K-means, and Louvain identified domains are 0.42, 0.24, and 0.33, respectively, suggesting that the SpaGCN results better agree with the manually curated layer structure reported in the original study.

**Figure 4.**
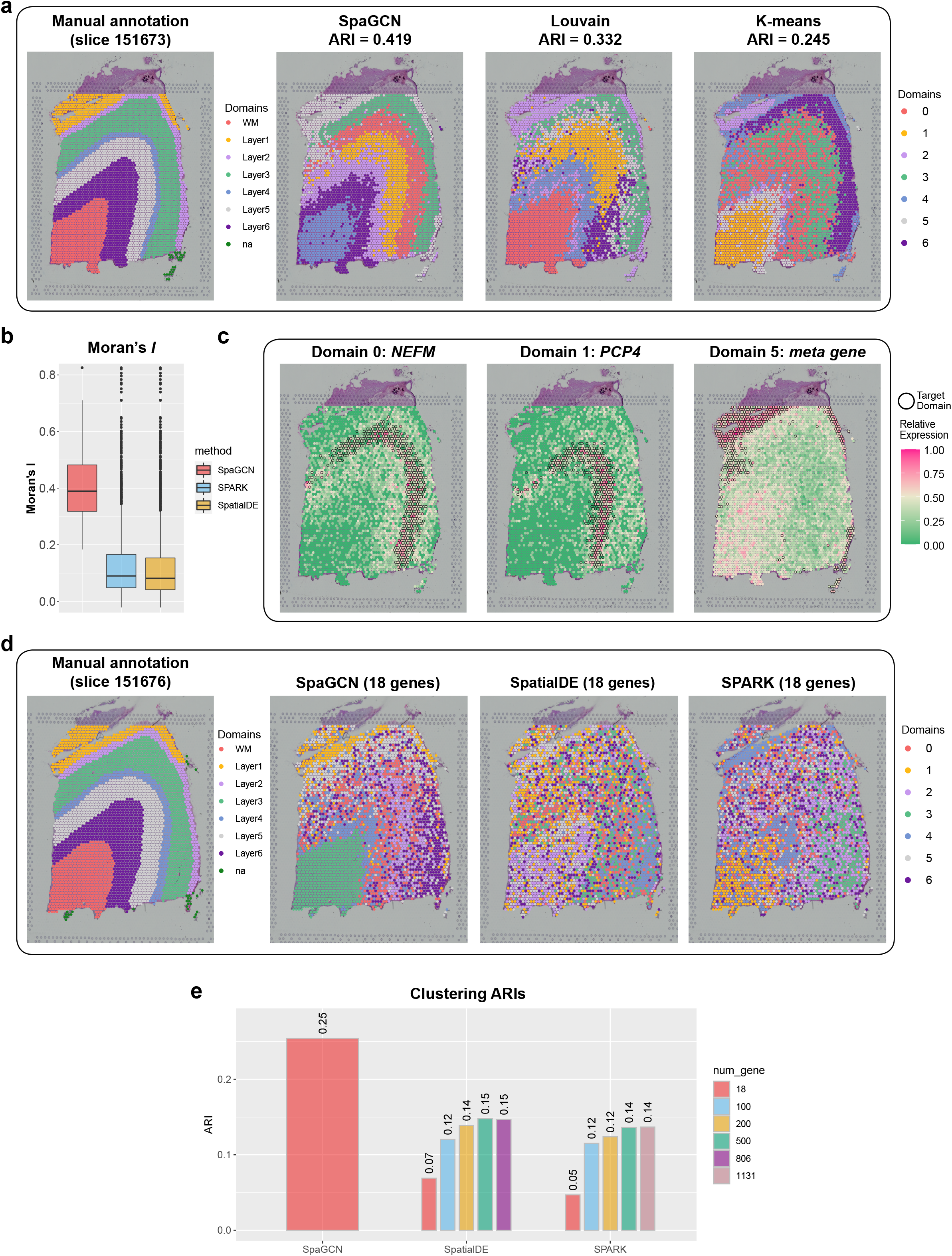
Spatial domains and SVGs detected in the LIBD human dorsolateral prefrontal cortex dataset. **a**, Manually annotated layer structure for slice 151673 from the original study^20^, and spatial domains detected by SpaGCN, Louvain’s method, and K-means clustering. **b**, Boxplot of Moran’s *I* values for SVGs detected by SpaGCN, SPARK, and SpatialDE for slice 151673. **c**, Spatial expression patterns of SVGs for domain 0 (*NEFM*) and domains 1 (*PCP4*), and a meta gene formed by *KRT19, MYL9, MBP, GFAP*, and *SNAP25* for domain 5 (*KRT19 + MYL9 – MBP + GFAP – SNAP25*). **d**, Manually annotated layer structure for slice 151676 from the original study^20^, and K-means clustering results for slice 151676 using 18 genes selected by SpaGCN, SPARK, and SpatialDE. For SpaGCN, we selected the following genes, domain 0 (*NEFL, NEFM*), domain 1 (*PCP4, TMSB10 + PCP4 – KRT19*), domain 2 (*CCK + KRT17 – MT-ND1, CPLX2 + KRT17 – MT-ND2*), domain 3 (*CAMK2N1, ENC1*), domain 4 (*MBP, FTL*), domain 5 (*KRT19 + MYL9 – MBP + GFAP – PLP1, KRT8 + MYL9 – MBP + GFAP – PLP1*), and domain 6 (*GFAP, MBP*), resulting in 18 unique genes in total. For SPARK and SpatialDE, the 18 SVGs with the smallest FDR-adjusted p-value or q-value were randomly selected. **e**, ARIs between manually annotated layers and K-means’ clustering using SVGs selected by different methods. For SpaGCN, we only used the selected SVGs and meta genes, with 18 genes involved in total while for SPARK and SpatialDE, we used top 18, 100, 200, 500 and all SVGs with the identical smallest FDR-adjusted p-value or q-value.

To further validate the identified spatial domains, we then detected SVGs. In total, SpaGCN detected 61 SVGs, with 53 of them specific to domain 4, which corresponds to the white matter region (Supplementary Fig. 14). Patterns of SVGs for other domains are not very clear. These results indicate that gene expression profiles of spots from white matter are distinct from spots in the neuronal layers, while gene expression differences among the six neuronal layers are much smaller and more difficult to distinguish using individual marker genes. SVGs detected by SPARK and SpatialDE also suffered from the same problem. SPARK detected 3,187 SVGs with 1,131 of them having FDR-adjusted p-values equal to 0, most of which only marked the white matter region. We also found that the SVGs detected by SPARK lack domain specificity (Supplementary Fig. 15). SpatialDE detected 3,654 SVGs with 806 of them having q-values equal to 0, but these genes do not necessarily show better spatial pattern than genes with larger q-values (Supplementary Fig. 16). Although SPARK and SpatialDE detected much larger numbers of SVGs than SpaGCN (Supplementary Fig. 17), the genes detected by these two methods lack ability to distinguish different degrees of spatial variability in expression as their p-value and q-value distributions are highly skewed toward 0 (Supplementary Fig. 18). Fig. 4b shows that the Moran’s *I* values for SpaGCN detected SVGs are significantly higher than those detected by SpatialDE and SPARK (median of 0.39 for SpaGCN against 0.09 for SPARK and 0.08 for SpatialDE). For 3 out of the 6 neuronal layers, SpaGCN detected a single SVG to mark that region (Fig. 4c). For example, *NEFM* is enriched in domain 0 (layer 3) and *PCP4* is enriched in domain 1 (layer 4). Although it is difficult to identify single genes to mark the other neuronal layers, SpaGCN was able to find layer-specific meta genes. As shown in Fig. 4c, the meta gene formed by *KRT19, MYL9, MBP, GFAP*, and *SNAP25* for domain 5 is specific to layer 1. Since layer 1 only has few spots, it is difficult to find a highly enriched gene. However, by adding depleted genes like *MBP* and SNAP25, the expression pattern in this region is strengthened. Furthermore, the SVGs and meta genes detected by SpaGCN are transferrable to slice 151676 obtained from the same study (Supplementary Fig. 19 and Supplementary Fig. 20).

To show the SVGs and meta genes detected by SpaGCN are useful for downstream analysis, we performed K-means clustering on slice 151676 using SVGs and meta genes detected from slice 151673 by SpaGCN. Specifically, we selected 2 SVGs or meta genes detected by SpaGCN for each spatial domain, resulting in 14 features (18 unique genes involved in total) used in K-means clustering. Comparing with manually curated layer assignment reported in the original study, this clustering analysis had an ARI of 0.25 (Fig. 4d). We performed similar clustering analysis using SVGs detected by SpatialDE and SPARK. When only using their top 18 SVGs, the ARI is only 0.07 for SpatialDE and 0.05 for SPARK. Even when using the 806 most significant SpatialDE detected SVGs, the ARI is only 0.14. When using the 1,114 most significant SPARK detected SVGs, the ARI is 0.15 (Fig. 4e). The ARIs of both SpatialDE and SPARK are much lower than SpaGCN, even though both used many more SVGs than SpaGCN, which further confirmed the lack of spatial expression specificity for genes detected by these methods.

### Application to human primary pancreatic cancer tissue

We also analyzed a human primary pancreatic cancer tissue dataset^5^, which includes 224 spots and 16,448 genes across 3 manually annotated sections, to show SpaGCN’s ability in detecting tumorous regions. The original study identified and annotated the cancer region on the histology image. However, the cancer region detected by their clustering method based on gene expression information alone did not closely match the pathologist annotated cancer region (Fig. 5a). Since the cancer region in the histology image is darker in color than non-cancer regions, it is informative for clustering. To give histology information higher weight, we increased the scaling parameter s in SpaGCN from 1 to 2 when calculating distance between each spot pair. This step ensured that spots in the same dark region in the histology are more likely to be clustered together. Fig. 5a shows that domain 2 detected by SpaGCN has a better correspondence to the cancer region than clusters reported in the original study. In total, SpaGCN detected 12 SVGs, with 3, 8, and 1 SVGs for domains 0, 1, and 2, respectively (Fig. 5b; Supplementary Fig. 21). Furthermore, a meta gene using *KRT17, MMP11*, and *SERPINA1* marked the cancer region better than the originally identified SVG *KRT17* (Fig. 5c). *KRT17* functions as a tumor promoter and regulates proliferation in pancreatic cancer^21^, and *MMP11* has been found to be a prognostic biomarker for pancreatic cancer^22^. Our identification of *KRT17* and *MMP11* as the two positive genes for the cancer region agree well with pancreatic cancer biology. SPARK and SpatialDE detected 203 and 163 SVGs, respectively (Supplementary Fig. 22). However, the Moran’s *I* values for their SVGs are much lower than those detected by SpaGCN, suggesting their lack of spatial expression patterns (Fig. 5d).

**Figure 5.**
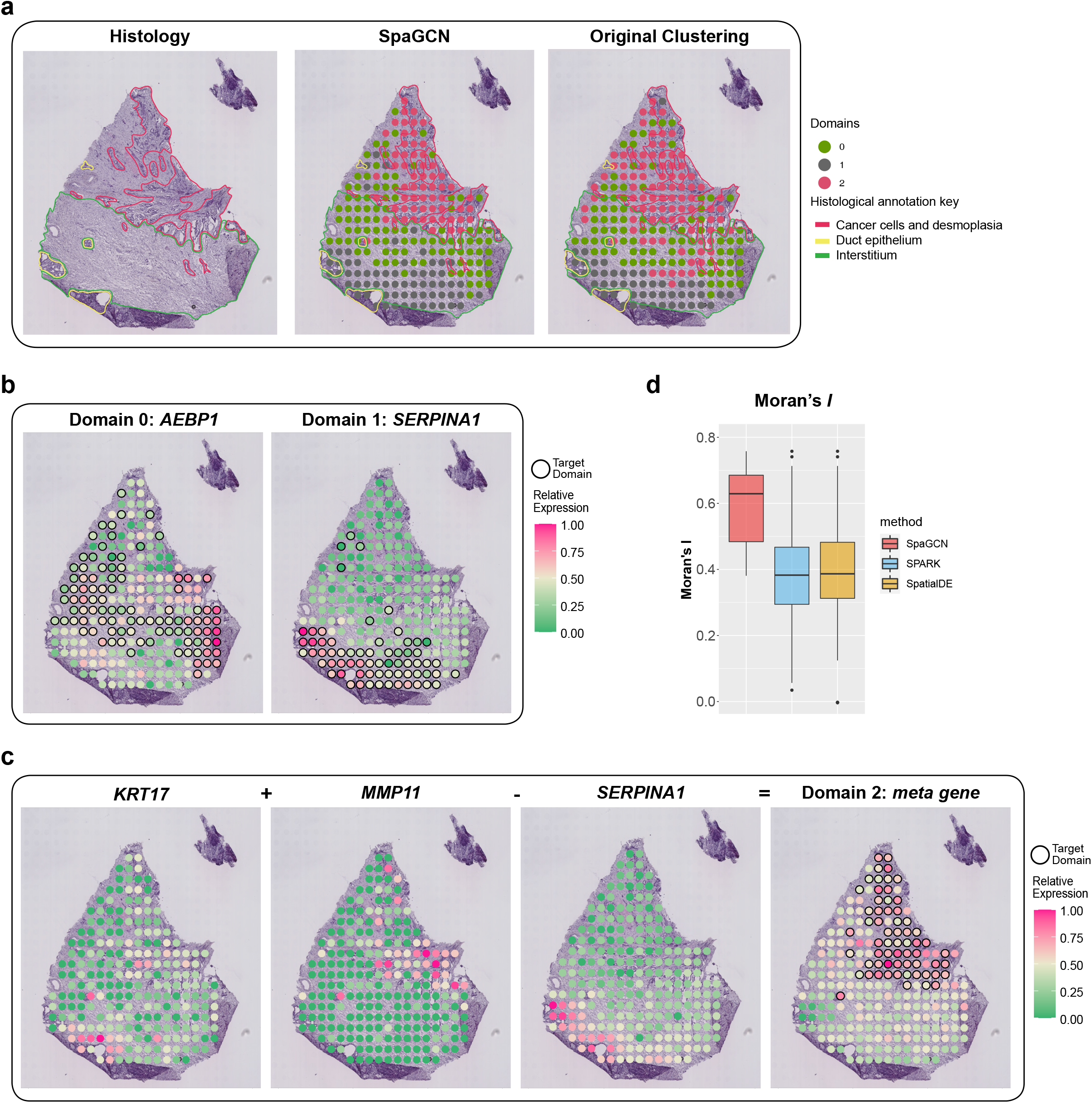
Spatial domains and SVGs detected in the human primary pancreatic cancer tissue dataset. **a**, Histology image of the tissue section with manually annotated regions from the original study^5^, spatial domains detected by SpaGCN, and clustering results from the original study. **b**, Spatial expression pattern of SVGs detected by SpaGCN for domain 0 (*AEBP1*) and domain 1 (*SERPINA1*). **c**, Spatial expression patterns of genes *KRT17, MMP11, SERPINA1*, which form the specific marker meta gene for domain 2 (*KRT17 + MMP11 - SERPINA1*). **d**, Boxplot of Moran’s *I* values for SVGs detected by SpaGCN, SPARK, and SpatialDE.

### Application to MERFISH mouse hypothalamus data

Next, we show that SpaGCN can also be applied to smFISH-based data. To this end, we analyzed a MERFISH dataset generated from the preoptic region of hypothalamus in mouse brain^2^, which includes 5,665 cells and 161 genes. One important difference between MERFISH and sequencing-based spatial transcriptomics data is that the captured tissue area is much smaller and less genes are measured, making it difficult to detect spatial domains since the cells within such a small area are more similar to each other. Thus, when utilizing these types of data, we suggest increasing the contribution of neighboring cells when calculating the weighted gene expression of each cell. Using this approach, SpaGCN detected spatial domains that agreed well with the annotated hypothalamic nuclei (Fig. 6a), with domain 2 corresponding to ACA, domain 3 corresponding to PS, and domain 7 corresponding to MnPo. By contrast, the domains identified from the Hidden Markov Random Field (HMRF) approach showed little overlap with the hypothalamic region annotation. Using SpaGCN, we further detected 19 SVGs including *DGKK, ERMN*, and *SLN* that showed enriched expression patterns for domains 2, 3, and 7 (Fig. 6b; Supplementary Fig. 23).

**Figure 6.**
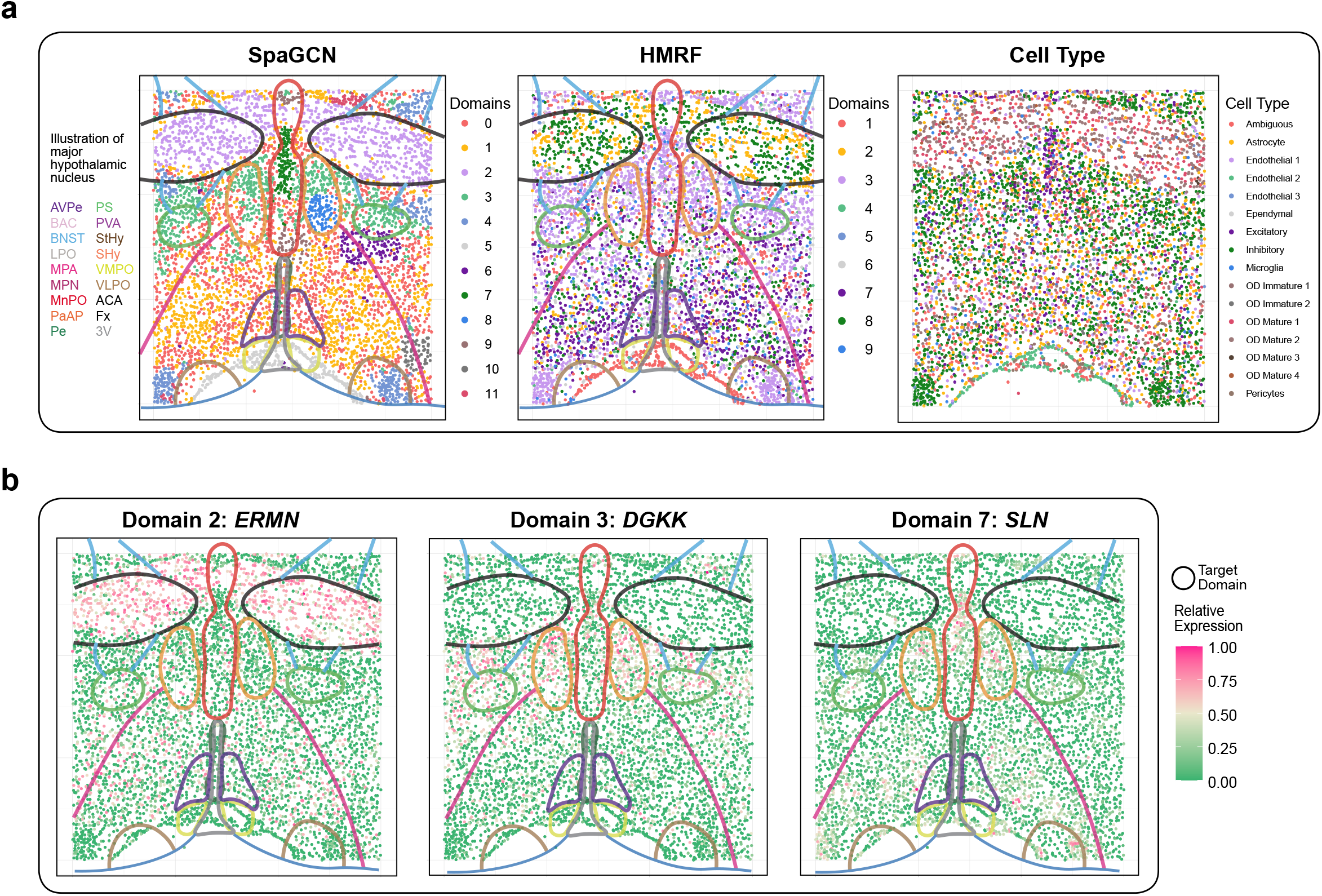
Spatial domains and SVGs detected in the MERFISH mouse brain hypothalamus dataset. **a**, Spatial domains detected by SpaGCN and the HMRF method overlayed with annotated hypothalamic nuclei from the original study^2^, and cell type distribution from the original study. **d**, Spatial expression patterns of SVGs detected by SpaGCN for domain 2 (*ERMN*), domain 3 (*DGKK*), and domain 7 (*SLN*).

## Discussion

Identification of spatial domains and detection of SVGs are important steps in spatial transcriptomics data analysis. In this paper, we presented SpaGCN, a graph convolutional network-based approach that integrates gene expression, spatial location, and histology to model spatial dependency of gene expression for clustering analysis of spatial domains and identification of domain enriched SVGs or meta genes. Through the use of a convolutional layer in an undirected weighted graph, SpaGCN aggregates gene expression of each spot from its neighboring spots, which enables the identification of spatial domains with coherent gene expression and histology. The subsequent domain guided differential expression analysis also enables the detection of SVGs or meta genes with enriched expression patterns in the identified domains. SpaGCN has been extensively tested on datasets from different species, regions, and tissues generated using both sequencing- and smFISH-based techniques. The results consistently showed that SpaGCN can identify spatial domains with coherent gene expression and histology and detect SVGs and meta genes that have much clearer spatial expression patterns and biological interpretations than genes detected by SPARK and SpatialDE. Additionally, the SpaGCN detected SVGs and meta genes are transferrable and can be utilized for downstream analyses in independent tissue sections.

The spatial domain detection step in SpaGCN is flexible. For datasets with clear layer structure in histology image, such as the mouse posterior brain data and human primary pancreatic cancer data, higher weight can be given to histology by increasing the scaling parameter s in SpaGCN when calculating distance between each spot pair, which results in spatial domains that are more similar to the anatomy-based taxonomy in the histology image. Another important scaling parameter in SpaGCN is the characteristic length scale *l*, which controls the relative contribution from other spots when aggregating gene expression. By varying *l*, users can get spatial domain separations with different patterns in which a higher *l* will result in spatial domains with higher contiguity.

The SVG detection procedure in SpaGCN is also flexible. While we mainly demonstrated SVG detection for a single domain, SpaGCN also allows users to combine multiple domains as one target domain or specify which neighboring domains to be included in DE analysis. Additionally, SpaGCN allows the users to customize SVG filtering criteria based on p-value and three additional metrics, i.e., in-fraction, in/out fraction ratio, and fold change, to select SVGs. The resulting genes can be ranked by any of these metrics to select SVGs with desired spatial expression patterns.

SpaGCN is computationally fast and memory efficient. To showcase the computational advantage of SpaGCN, we recorded its run time and memory usage for the mouse posterior brain data and compared with SPARK and SpatialDE. All analyses were conducted on Mac OS 10.13.6 with single Intel® Core(TM) i5-8259U CPU @2.30GHz and 16GB memory. As shown in Supplementary Fig. 24, SpaGCN completed spatial domain and SVG detection in less than one minute, whereas the computing time is ~13 minutes for SpatialDE and more than 18 hours for SPARK. Furthermore, SpaGCN only required 1.3 GB of memory, whereas SpatialDE and SPARK required more than 3.1 GB and 7.2 GB of memory, respectively. With the increasing popularity of spatial transcriptomics in biomedical research, we expect SpaGCN will be an attractive tool for large-scale spatial transcriptomics data analysis. Results from SpaGCN will enable researchers to accurately identify spatial domains and SVGs in their studies.

## Acknowledgements

This work was supported by the following grants: R01GM125301, R01EY030192, R01EY031209 R01HL113147, and R01HL150359 (to M.L.), and P01AG066597 (to D.J.I. and E.B.L.). We thank Reuben Moncada and Itai Yanai for sharing the human pancreatic cancer histology image data.

## Author contributions

This study was conceived of and led by M.L.. J.H. designed the model and algorithm. J.H. implemented the SpaGCN software and led the data analysis with input from M.L., X.L., K.C., A.S., D.I., E.L., and R.T.S.. J.H. and M.L. wrote the paper with feedback from all other coauthors.

## Methods

### Data preprocessing

SpaGCN takes spatial gene expression and histology image data (when available) as input. The spatial gene expression data are stored in an *N × D* matrix of unique molecular identifier (UMI) counts with *N* samples and *D* genes, along with the (*x, y*) 2-dimensional spatial coordinates of each sample. In sequencing-based data, each sample represents a spot containing multiple cells, whereas in single-molecule fluorescence *in situ* hybridization (smFISH)-based data, each sample represents a single cell. For simplicity, we will use ‘spot’ to refer to a sample, as most of the data analyzed in this paper are sequencing based. Genes expressed in less than three spots are eliminated. The gene expression values in each spot are normalized such that the unique molecular identifier (UMI) count for each gene is divided by the total UMI count across all genes in a given spot, multiplied by 10,000, and then transformed to a natural log scale.

### Conversion of spatial transcriptomics data into graph-structured data

After preprocessing, SpaGCN converts the gene expression and histology image data into a weighted undirected graph, *G*(*V, E*). In this graph, each vertex *v ∈ V* represents a spot and every two vertices in *V* are connected via an edge with a specified weight. We focus our analysis on spatial transcriptomics data with histology information, but the method can be easily adapted to analyze smFISH-based data, for which histology information is not available.

#### Calculation of distance between two vertices

The distance between any two vertices *u* and *v* in the graph reflects the relative similarity of the two corresponding spots. This distance is determined by two factors: 1) the physical locations of spots *u* and *v* in the tissue slice, and 2) the corresponding histology information of these two spots. Although some spots are physically close to each other in the tissue, the histology image may reveal that they belong to different tissue layers. Therefore, SpaGCN considers two spots to be close if and only if 1) the two spots are physically close, and 2) they have similar pixel features as shown in the histology image. To define a distance metric considering both aspects, SpaGCN extends the 2-dimensional space in the tissue slice into a 3-dimensional space that incorporates histology information. For spot *v*, its physical location in the tissue slice is represented by 2-dimensional coordinates (*x_v_, y_v_*). To determine the corresponding pixel in the histology image for spot *v*, SpaGCN maps spot *v* to the histology image according to its pixel coordinates (*x_pv_, y_pv_*). Instead of using the color of the pixel at (*x_pv_, y_pv_*), SpaGCN draws a square centered on (*x_pv_, y_pv_*) containing 50 × 50 pixels and calculates the mean color value for the RGB channels, (*r_v_, g_v_, b_v_*), of all pixels that fall in the square. This step smooths the color value and ensures that the color is not dominated by a single pixel. To derive a single value to represent the histology image features, SpaGCN uses a weighted sum of the RGB values as follows,

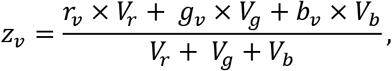

where *V_r_* = Variance(*r_v_*), *V_g_* = Variance(*g_v_*), and *V_b_* = Variance(*b_v_*) for all *v ∈ V*. In this transformation, higher weight is given to the channel with larger variance so that this combined value *z_v_* captures an accurate representation of the patterns in the histology image.

Next, SpaGCN rescales *z_v_* as

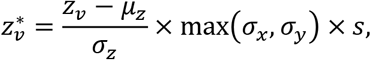

where *μ_z_* is the mean of *z_v_, σ_x_, σ_y_, σ_z_* are the standard deviations of *x_v_*, *y_v_* and *z_v_*, respectively, for *v ∈ V*, and *s* is a scaling factor. In our analysis, *s* is usually set at 1 to make sure that 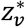 has the same scale variance as *x_v_* and *y_v_*, and we set s to a value larger than 1 when the goal is to increase the weight of histology. The coordinates of spot *v* are set to be 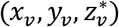 in the extended 3-dimensional space. Finally, the Euclidean distance between every two spots *u* and *v* is calculated as

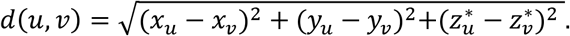

#### Calculation of weight for each edge and construction of graph

The weight of each edge (*u, v*) measures the degree of relatedness between spots *u* and *v* and is negatively associated with their distance. The graph structure *G* is stored in an *N × N* adjacency matrix ***A*** = [*w*(*u, v*)], where the edge weight between spot *u* and spot *v* and is defined as

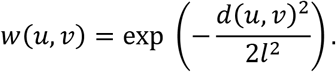

The hyperparameter *l*, also known as the characteristic length scale, determines how rapidly the weight decays as a function of distance. A similar function has been employed in SpatialDE^13^. Let ***I*** denote the identity matrix. For spot *v*, the corresponding row sum of ***A – I***, denoted by *a_v_*, can be interpreted as the relative contribution of other spots to its gene expression. We choose the value of *l* such that the average of *a_v_* across all spots is equal to a pre-specified value, e.g. 0.5.

### Graph convolutional layer

SpaGCN reduces the dimension of the preprocessed gene expression matrix using principal component analysis (PCA). The top 50 principal components are used as input, which work well for all datasets analyzed in this paper. Next, utilizing the power of a graph convolutional network, SpaGCN concatenates the gene expression information and edge weights in *G* to cluster the nodes. Following Kipf and Welling^23^, the graph convolutional layer can be written as

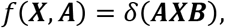

where ***X*** is the *N* × 50 embedding matrix obtained from PCA, ***B*** is a 50 × 50 matrix representing filter parameters of the convolutional layer, and *δ*(·) is a non-linear activation function such as ReLU. The graph convolutional layer ensures that a corresponding row of parameters in ***B*** will control the aggregation of neighborhood information for each feature in ***X***, thus offering the flexibility of feature specific aggregation of information provided by neighboring spots. The filter parameters in ***B*** are shared across all vertices in the graph and are automatically updated during an iterative training progress. Through graph convolution, SpaGCN has aggregated the gene expression information according to the edge weights specified in *G*. The output of this layer is an aggregated matrix that includes information on gene expression, spatial location, and histology. The graph convolutional layer was implemented based on Kipf and Welling^23^, where the backpropagation is operated via a localized first-order approximation of spectral graph convolution.

### Spatial domain identification by clustering

Next, based on the output from the above graph convolutional layer, SpaGCN employs an unsupervised clustering algorithm to iteratively cluster the spots into different spatial domains^15^. Each cluster identified from this analysis is considered to be a spatial domain, which contains spots that are coherent in gene expression and histology. To initialize cluster centroids, we use Louvain’s method^7^ on the aggregated output matrix from the graph convolutional layer. If the number of domains in the tissue is known, the resolution parameter in Louvain will be set to generate the same number of spatial domains. Otherwise, we vary the resolution parameter from 0.2 to 1.0 and select the resolution that gives the highest Silhouette score^24^.

To update the cluster assignments iteratively, we define a metric to measure the distance from a spot to a cluster centroid using the Student’s *t*-distribution as a kernel. The distance between the embedded point *h_i_* for spot *i* and centroid *μ_j_* for cluster *j*

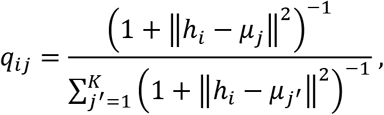

can be interpreted as the probability of assigning cell *i* to cluster *j*.

Next, we iteratively refine the clusters by defining an auxiliary target distribution *P* based on *q_ij_*

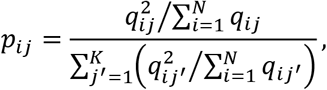

which upweights spots assigned with high confidence, and normalizes the contribution of each centroid to the overall loss function to prevent large clusters from distorting the hidden feature space. Now that we have the soft assignment *q_ij_* and the auxiliary distribution *p_ij_*, we can define the objective function as a Kullback-Leibler (KL) divergence loss,

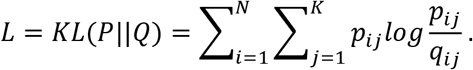

The network parameters and cluster centroids are simultaneously optimized by minimizing *L* using stochastic gradient descent with momentum. This unsupervised iterative clustering algorithm has been previously utilized for scRNA-seq analysis and showed superior performance over Louvain’s method^25,26^.

### Detection of spatially variable genes

We are interested in detecting spatially variable genes (SVGs) that are enriched in each spatial domain. We note that some genes may be expressed in multiple but disconnected domains. Although they are not uniquely expressed in a particular domain, these genes are still useful for understanding spatial variation of gene expression and can be used to form meta genes that are uniquely expressed in a specific domain. Therefore, rather than doing differential expression (DE) analysis using spots from a target domain versus all other spots, we first select spots to form a neighboring set of the target domain. The goal is to detect genes that are highly expressed in the target domain but are not expressed or are expressed at low levels in the neighboring spots. To determine which spots should be considered as neighbors, we draw a circle with a prespecified radius around each spot in the target domain. All spots from non-target domains that reside in the circle are considered its neighbors. The radius is set such that all spots in the target domain have approximately 8 neighbors on average. Next, neighbors of all spots in the target domain are collected and form a neighboring set. For each non-target domain, if more than 50% (default) of its spots are in the neighboring set, this domain is then selected as a neighboring domain. This criterion is set to avoid the situation when a domain is selected as a neighboring domain, but only a small proportion of its spots are adjacent to the target domain.

After neighboring domains are determined, SpaGCN then performs DE analysis between spots in the target domain and the neighboring domain(s) using Wilcoxon rank-sum test. Genes with a false discovery rate (FDR) adjusted p-value <0.05 are selected as SVGs. To ensure only genes with enriched expression patterns in the target domain are selected, we further require a gene to meet the following three criteria: 1) the percentage of spots expressing the gene in the target domain, i.e., in-fraction, is >80%; 2) for each neighboring domain, the ratio of the percentages of spots expressing the gene in the target domain and the neighboring domain(s), i.e., in/out fraction ratio, is >1; and 3) the expression fold change between the target and neighboring domain(s) is >1.5. If a user is interested in finding SVGs for a particular combination of spatial domains, SpaGCN offers the option to do so.

### Detection of spatially variable meta genes

The spatial domain-specific DE analysis described above typically detects SVGs with enriched expression for the majority of the domains. For domains in which no such SVGs are detected, we aim to identify a set of genes that, when combined to form a meta gene, shows an enriched expression pattern in the given domain. To identify genes to form a meta gene, we employ a multi-step approach. First, we lower the thresholds for SVG filtering, e.g., change the minimum fold change threshold from 1.5 to 1.2, to identify genes showing weaker enriched expression pattern in the target domain. In the presence of multiple such weaker SVGs, we randomly select one of them as the base gene and denote it as *gene*_0_. Second, we aim to aggregate expression from other genes to the base gene to enhance the spatial pattern for the target domain. To achieve this goal, we first calculate the mean expression level of *gene*_0_ for spots in the target domain as *e*_0_. Then, all spots from non-target domains with *gene*_0_’s expression level higher than *e*_0_ are extracted to form a control group. Next, we perform DE analysis using spots from the target domain against spots in the control group using Wilcoxon rank-sum test. The gene with the smallest FDR-adjusted p-value and higher expression in the target domain is selected as *gene*_0+_. Similarly, we perform DE analysis using spots from the control group against those from the target domain and select a gene with the smallest FDR-adjusted p-value and higher expression in the control group as *gene*_0−_. The meta gene’s expression is calculated as

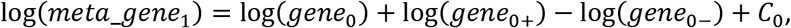

where *C*_0_ is a constant to make log(*meta_gene*_1_) non-negative. The log transformation is used to rescale expression and make the expression levels comparable across different genes. We have found that including negative genes can strengthen spatial expression pattern for domains that do not have enriched positive marker genes. This algorithm can be used iteratively to find additional genes to form an updated meta gene with a clearer spatial pattern for the target domain. For the (*t* + 1)^*th*^ iteration, the meta gene expression is calculated as

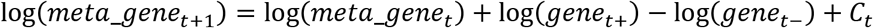

In the (*t* + 1)^*th*^ iteration, after adding *gene*_*t*+_ and subtracting *gene*_t−_, SpaGCN will select the (*t* + 1)^*th*^ control group based on *meta_gene*_*t*+1_. The size of the new control group, which is the number of spots not in the target domain but have higher expression of *meta_gene*_*t*+1_ than spots in the target domain, should be smaller than the size of the *t^th^* control group, to ensure that *meta_gene*_*t*+1_ has a clearer spatial pattern than *meta_gene_t_*. Also, *meta_gene*_*t*+1_ is expected to have a larger difference of mean expression between the target and control groups than *meta_gene_t_*. Therefore, at each iteration, SpaGCN checks whether both criteria are met, and the search of additional genes will stop otherwise. An illustration of this iterative meta gene search is shown in Supplementary Fig. 25.

### Evaluation of spatially variable genes using Moran’s *I* statistic

The Moran’s *I* statistic^16^ is a measure of spatial autocorrelation, which can be used to measure the degree of spatial variability in gene expression^27^. The Moran’s *I* value ranges from −1 to 1, where a value close to 1 indicates a clear spatial pattern, a value close to 0 indicates random spatial expression, and a value close to −1 indicates a chess board like pattern. To evaluate the spatial variability of a given gene, we calculate the Moran’s *I* using the following formula,

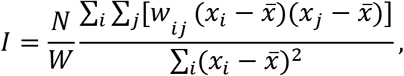

where *x_i_* and *x_j_* are gene expression of spots *i* and *j*, 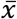 is the mean expression of the gene, *N* is the total number of spots, *w_ij_* is spatial weight between spots *i* and *j* calculated using the 2-dimensional spatial coordinates of the spots, and *W* is the sum of *w_ij_*. For each spot, we select the *k* nearest neighbors using spatial coordinates. Moran’s *I* statistic is robust to the choice of *k* and is set at 4 in our analysis. We assign *W_ij_* = 1 if spot *j* is in the nearest neighbors of spot *i*, and *w_ij_* = 0 otherwise.

### Detection of subclusters within a spatial domain

To better characterize heterogeneity within a spatial domain due to the influence of its neighborhood, SpaGCN can further detect sub-domains within each spatial domain by utilizing information from neighboring spots. SpaGCN draws a circle around each spot with a pre-specified radius, and all spots that reside in the circle are considered as neighbors of this spot. The value of the radius is set to ensure that every spot in the target domain have ten neighbors on average. Next, SpaGCN records the number of neighbors from different spatial domains for each spot and stores this information in a *T × K* matrix, where *T* is the number of spots in the target domain and *K* is the total number of spatial domains detected. The value for the *i^th^* row and *j^th^* column is the number of neighbors of spot *i* belonging to domain *j*. Next, this matrix is fed into a *K*-means classifier to detect sub-clusters. Differential expression analysis as described above can be performed to identify subcluster enriched genes.

### Data availability

We analyzed multiple spatial transcriptomics datasets. Publicly available data were acquired from the following websites or accession numbers: (1) mouse olfactory bulb (https://drive.google.com/drive/folders/1C4l3lBaYl7uuV2AA2o0WDzO_mkc_b0pv?usp=sharing); (2) mouse posterior brain (https://support.10xgenomics.com/spatial-gene-expression/datasets/1.0.0/V1_Mouse_Brain_Sagittal_Posterior); (3) LIBD human dorsolateral prefrontal cortex Dorsolateral pre-frontal cortex (http://research.libd.org/spatialLIBD/); (4) human primary pancreatic cancer data (GSE111672); (5) MERFISH mouse hypothalamus data (https://datadryad.org/stash/dataset/doi:10.5061/dryad.8t8s248). Details of the datasets analyzed in this paper were described in **Supplementary Table 1.**

### Software availability

An open-source implementation of the SpaGCN algorithm can be downloaded from https://github.com/jianhuupenn/SpaGCN

### Life sciences reporting summary

Further information on experimental design is available in the Life Sciences Reporting Summary.

