## Supplementary Materials for "Integrating gene expression, spatial location and histology to identify spatial domains and spatially variable genes by graph convolutional network"

***Correspondence:**

**Supplementary Table 1. Datasets analyzed in this paper.**

| Species | Tissue | Data Source | Dataset Dimensions | Protocol |
| --- | --- | --- | --- | --- |
| Mouse | Olfactory bulb | Ståhl *et al*. [1]  (https://drive.google.com/drive/folders/1C4l3lBaYl7uuV2AA2o0WDzO_mkc_b0pv?usp=sharing) | 262 spots  16,218 genes | Spatsial Transcriptomics |
| Mouse | Posterior brain (sagittal) | 10x Genomics  (https://support.10xgenomics.com/spatial-gene-expression/datasets/) | 3,353 spots  31,053 genes | 10X Visium |
| Human | Dorsolateral prefrontal cortex | Maynard *et al*. [2]  (http://research.libd.org/spatialLIBD/) | Sample 151673:  3,639 spots  33,538 genes  Sample 151676:  3,460 spots  33,538 genes | 10X Visium |
| Human | Primary pancreatic cancer tissue | Moncada *et al*. [3]  GSE111672 | 224 spots  16,448 genes | Spatial Transcriptomics |
| Mouse | Hypothalamus | Moffitt *et al*. [4]  GSE71585 | 5,665 cells  161 genes | MERFISH |

**Supplementary Table 2. Software compared with SpaGCN**.

| **Method** | **Version** | **URL** | **Reference** |
| --- | --- | --- | --- |
| SpatialDE | 1.1.3 | https://github.com/Teichlab/SpatialDE | [5] |
| SPARK | 1.0.2 | https://github.com/xzhoulab/SPARK | [6] |
| HMRF | 1.3.3 | https://bitbucket.org/qzhudfci/smfishhmrf-py/src/master/ | [7] |

**Supplementary Figure 1.** Gene expression patterns for 40 SVGs randomly selected from the 61 SVGs that were identified by SpaGCN in the mouse olfactory bulb dataset.

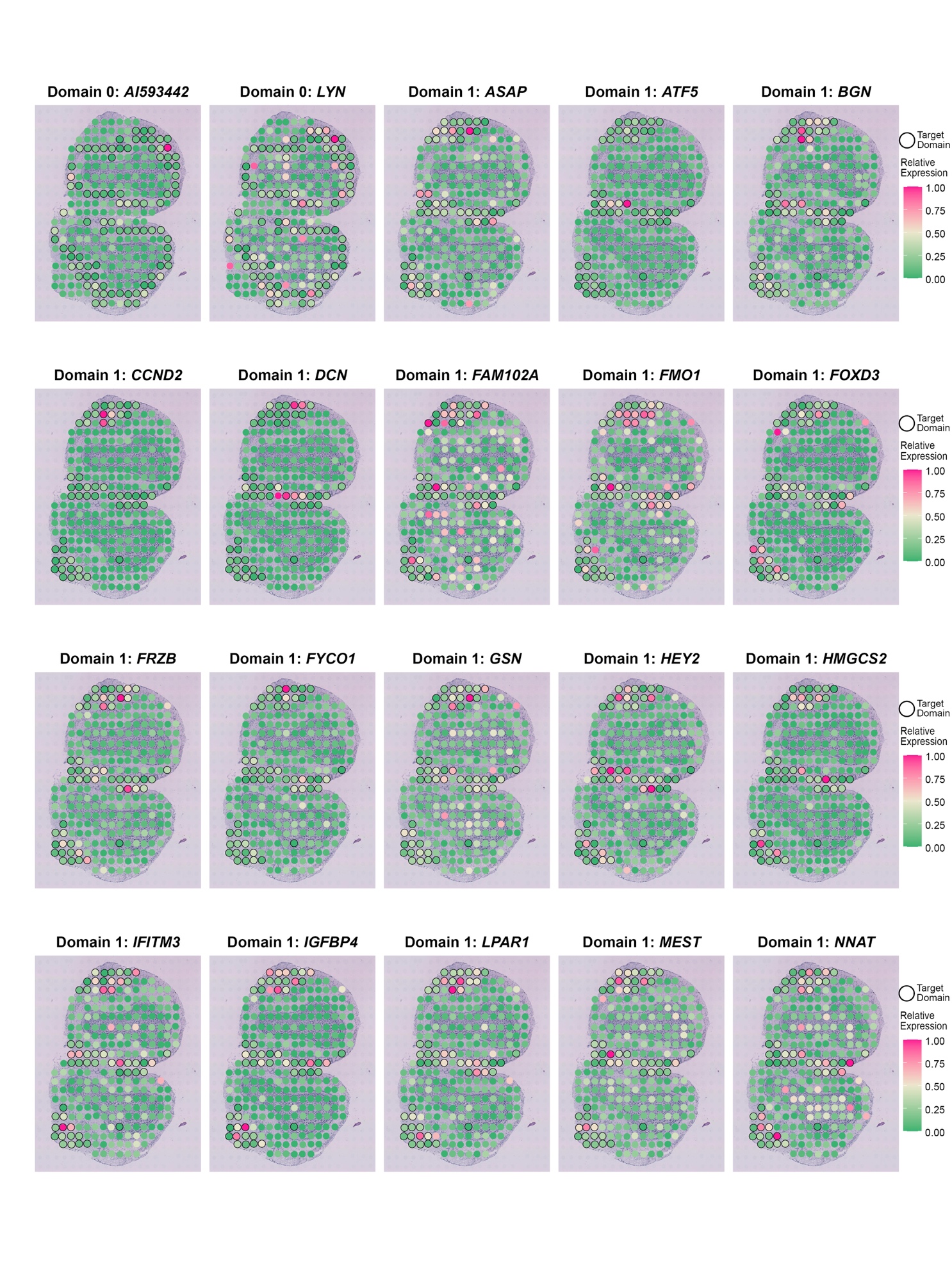

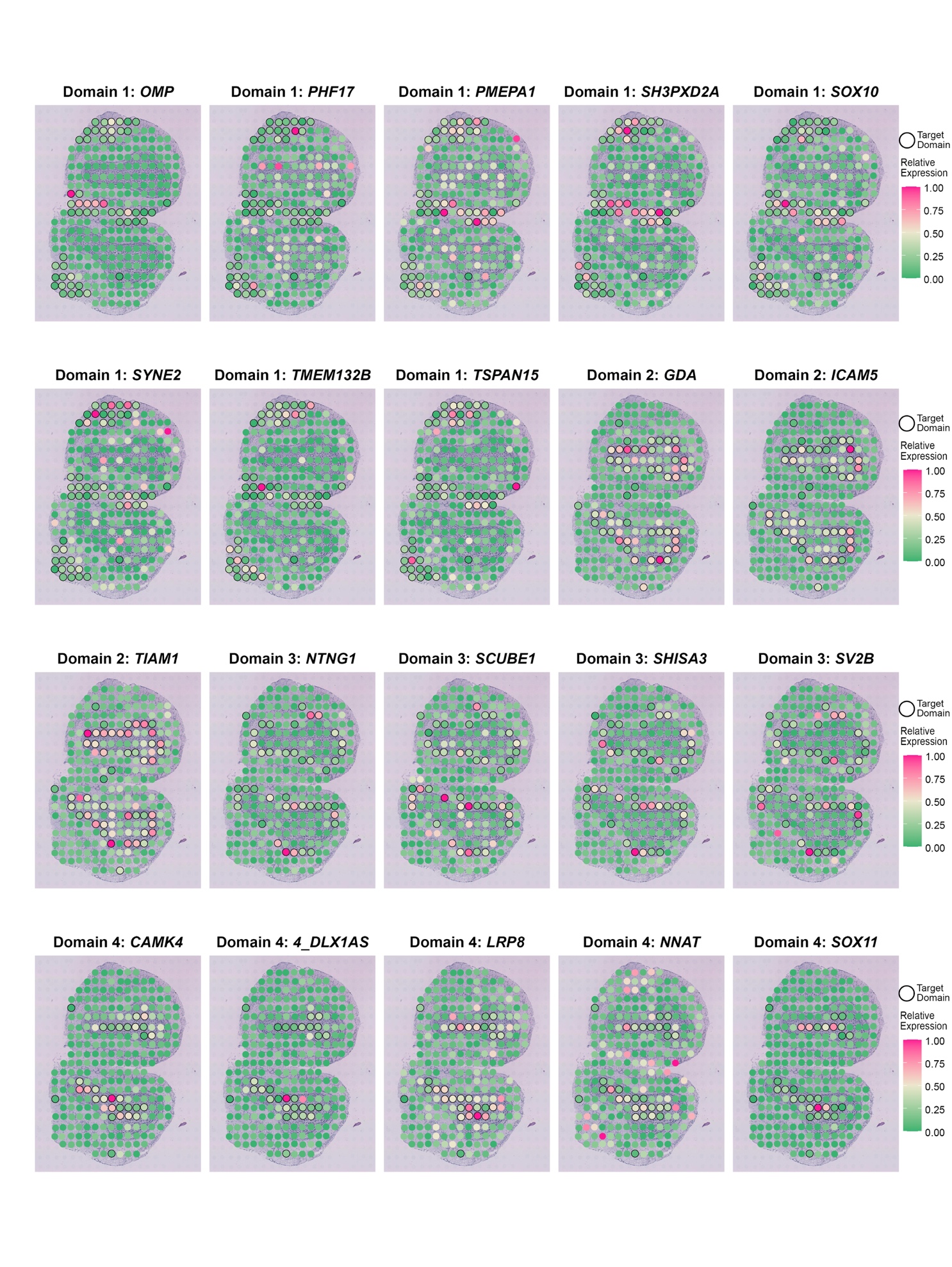

**Supplementary Figure 2.** Venn diagram for SVGs detected by SpaGCN, SpatialDE and SPARK in the mouse olfactory bulb dataset.

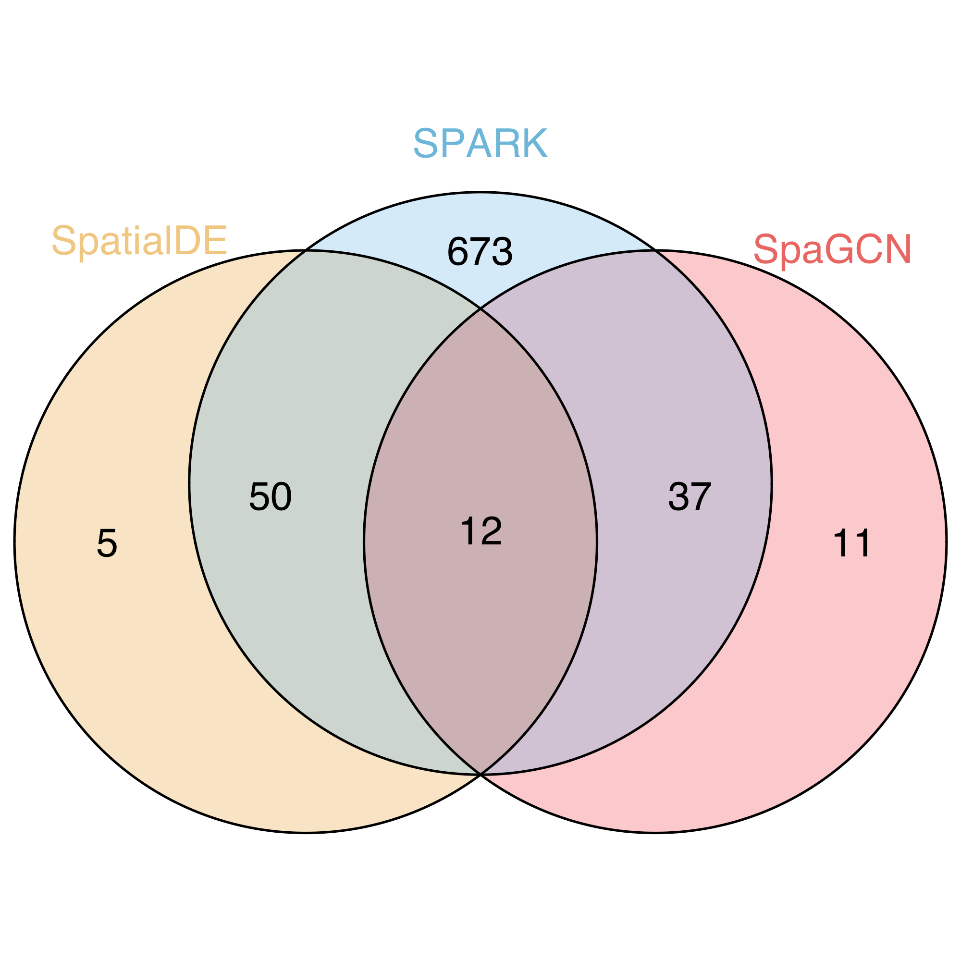

**Supplementary Figure 3.** Gene expression patterns for 20 SVGs randomly selected from the 55 SVGs that were identified by SpatialDE but not by SpaGCN in the mouse olfactory bulb dataset. These 55 SVGs have the smallest identical q-value 1.15e-09 from SpatialDE.

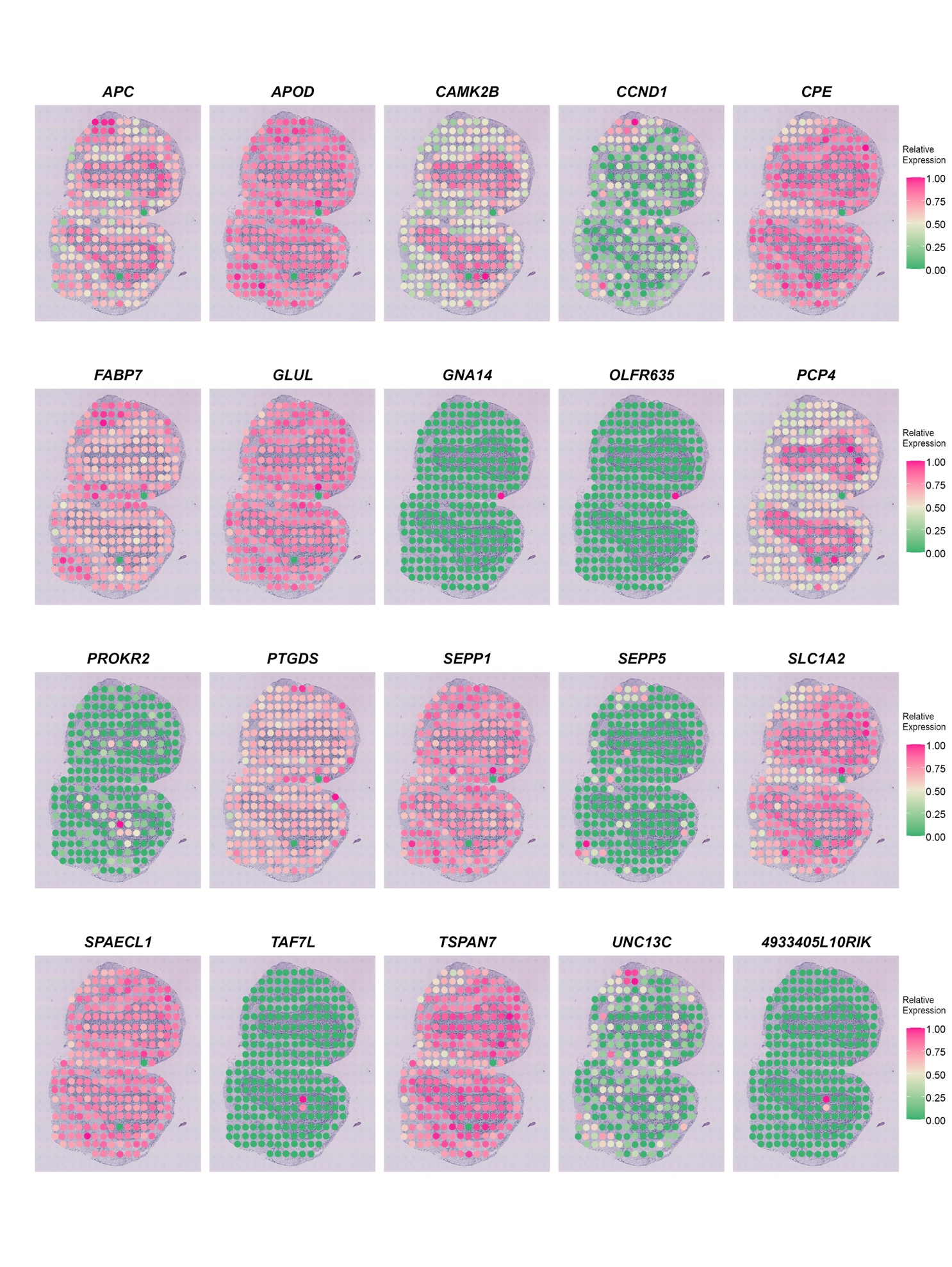

**Supplementary Figure 4.** Gene expression patterns for 14 SVGs with the smallest identical FDR-adjusted p-value 4.43e-13 detected by SPARK but not by SpaGCN in the mouse olfactory bulb dataset.

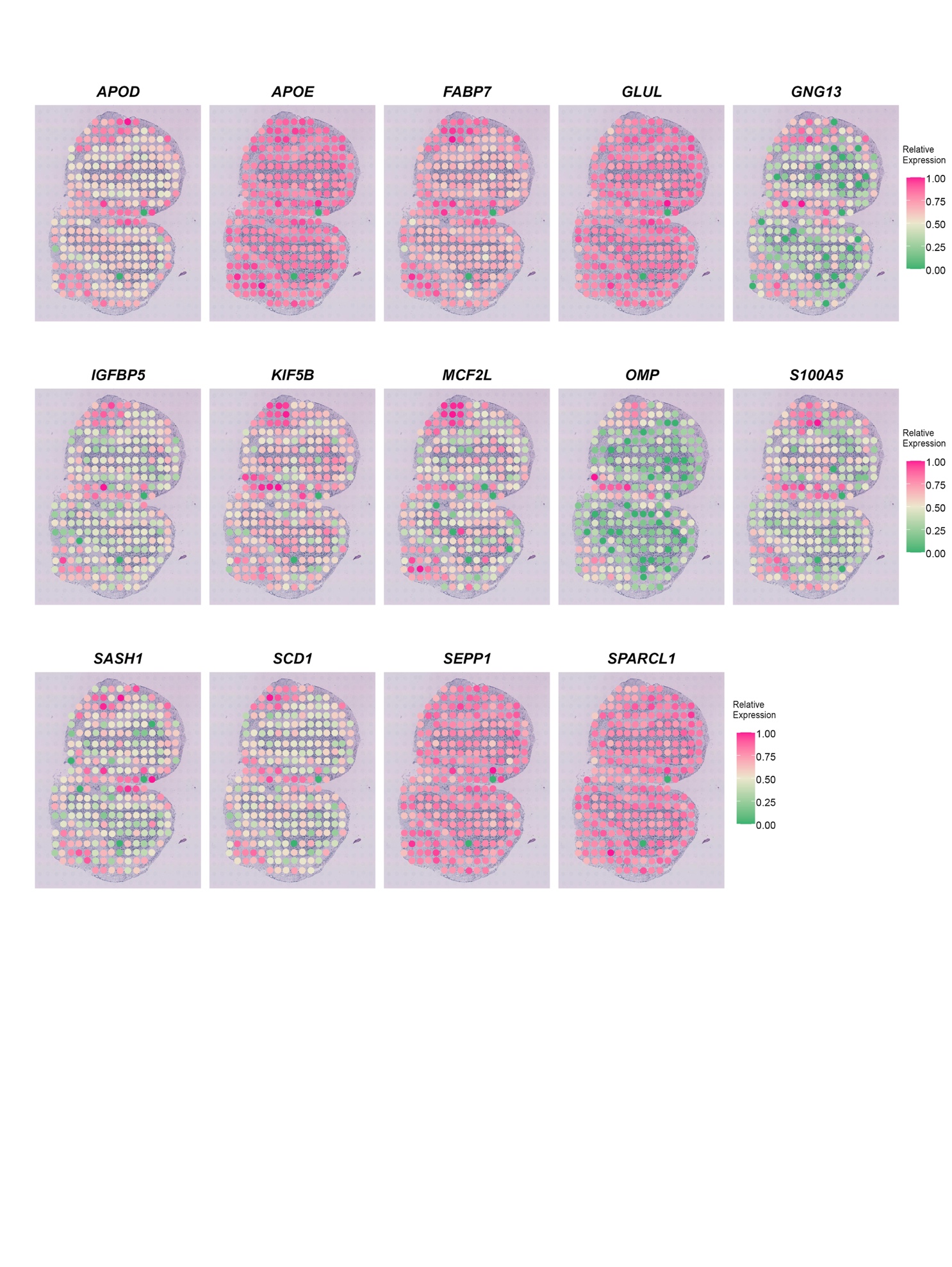

**Supplementary Figure 5.** P-value (FDR adjusted) distribution for SVGs detected by SPARK (772 genes total) and q-value distribution for SVGs detected by SpatialDE (67 genes total) in the mouse olfactory bulb dataset.

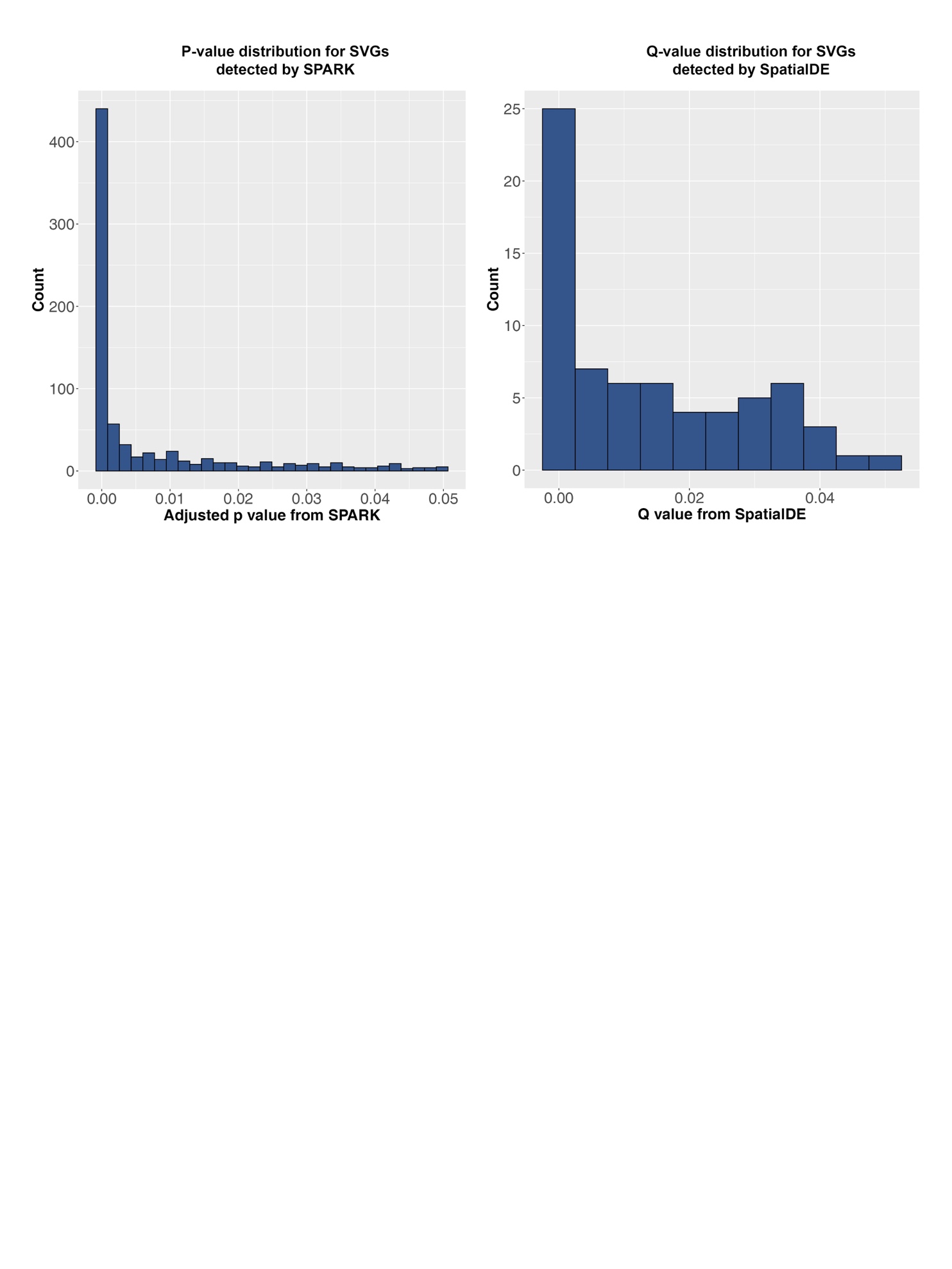

**Supplementary Figure 6.** SVGs detected by SPARK and SpatialDE that are expressed in multiple adjacent spatial domains in the mouse olfactory bulb dataset.

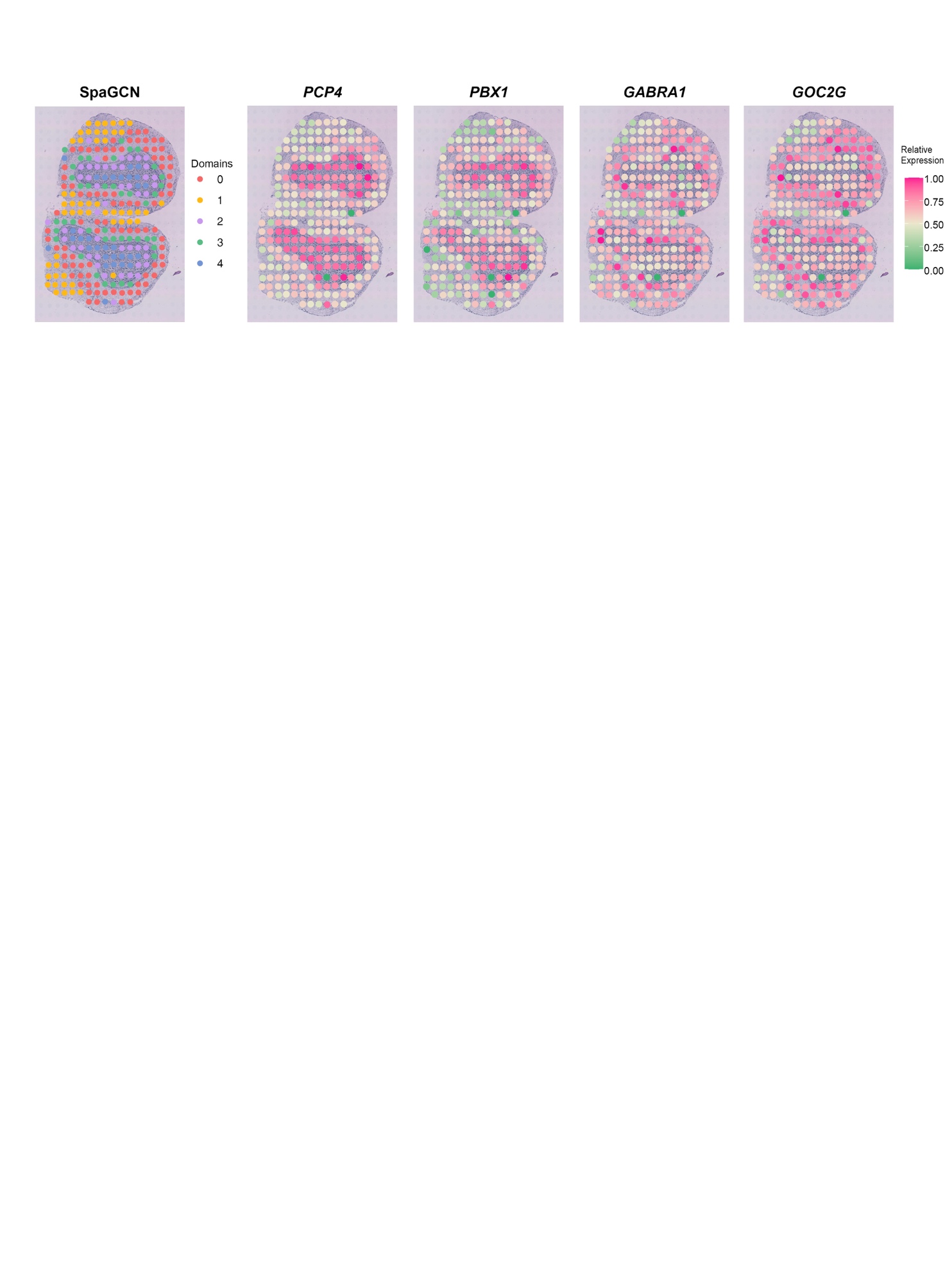

**Supplementary Figure 7.** Venn diagram for SVGs detected by SpaGCN, SpatialDE and SPARK in the mouse posterior brain dataset.

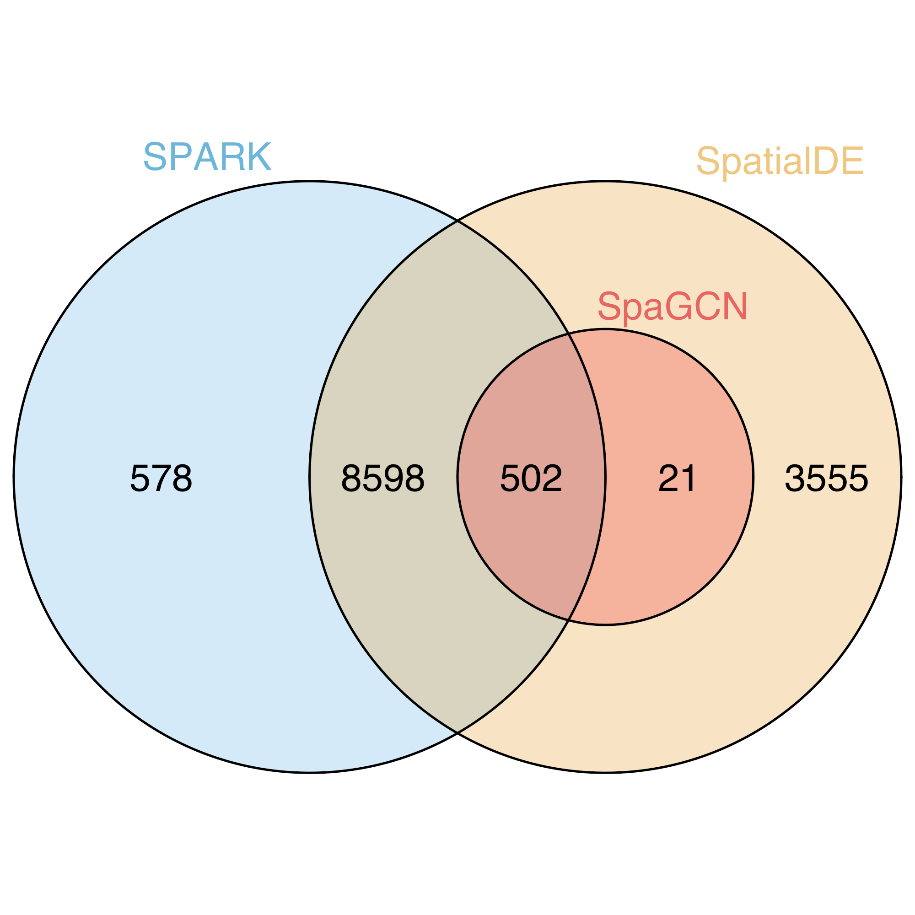

**Supplementary Figure 8.** SVGs detected by SPARK with smaller FDR-adjusted p-values do not necessarily show better spatial pattern than genes with larger p-values in the mouse posterior brain dataset. The p-values shown below are FDR-adjusted p-values.

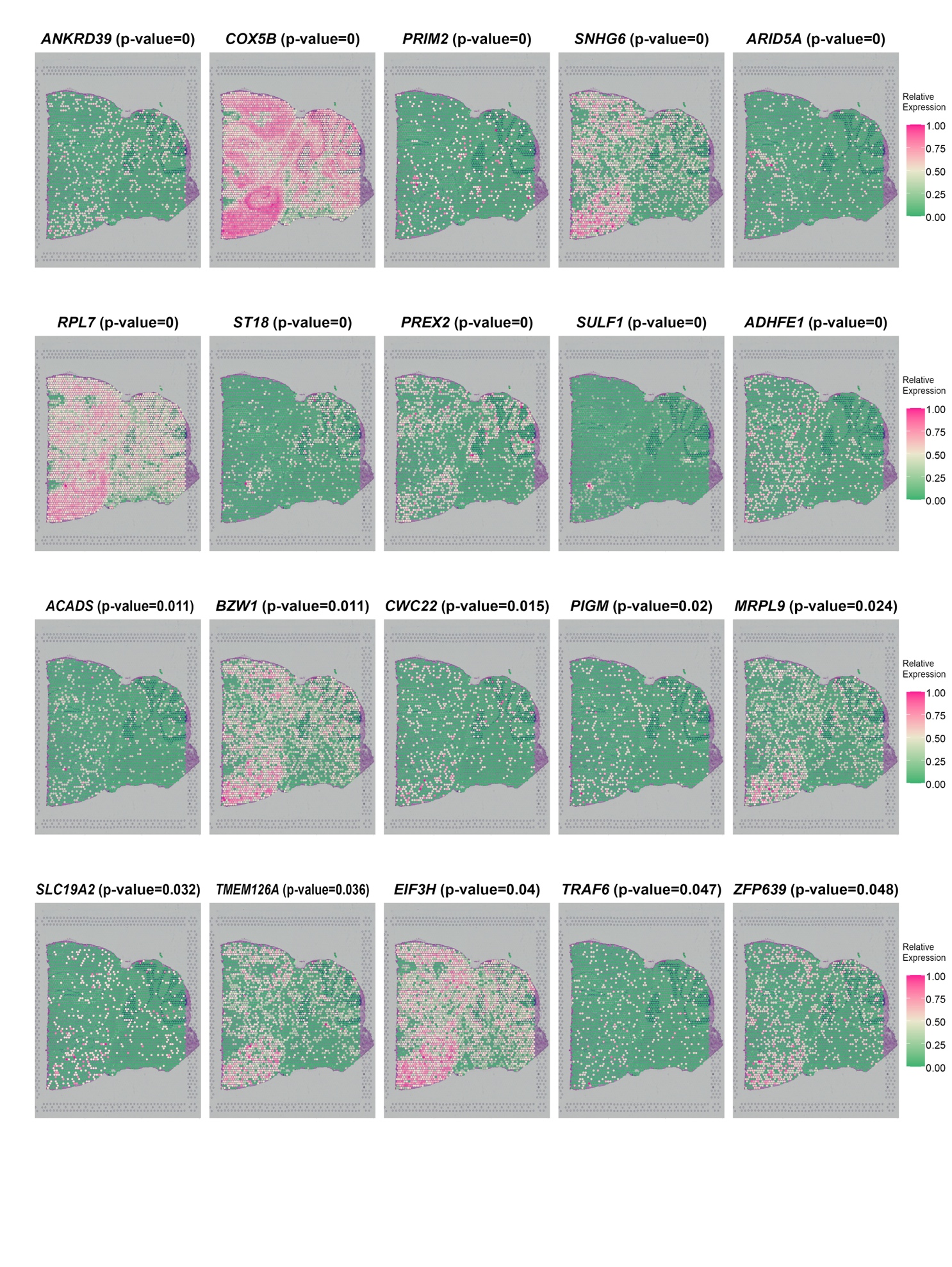

**Supplementary Figure 9.** SVGs detected by SpatialDE with smaller q-values do not necessarily show better spatial pattern than genes with larger q-values in the mouse posterior brain dataset.

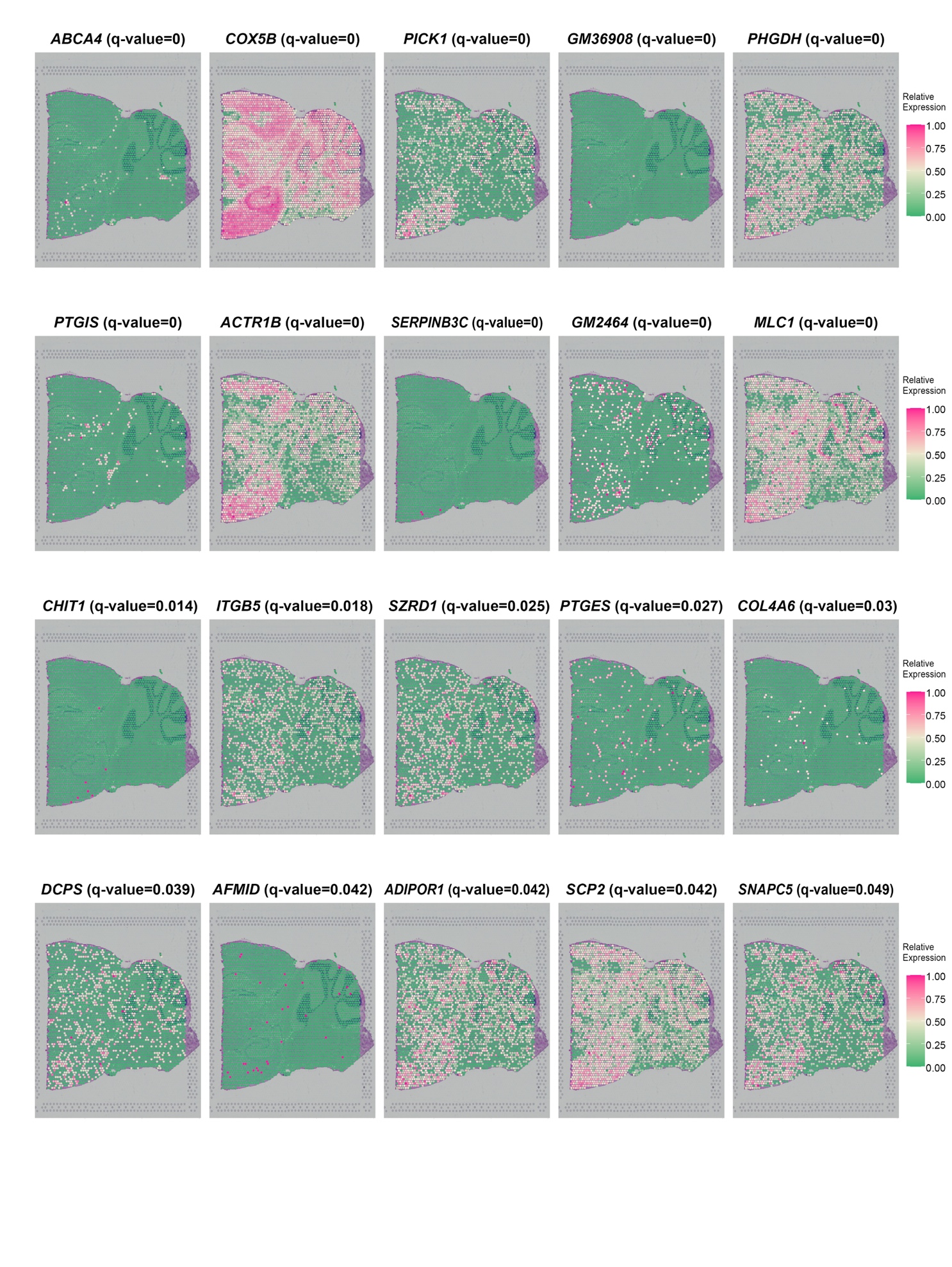

**Supplementary Figure 10.** P-value (FDR adjusted) distribution for SVGs detected by SPARK (9,678 genes in total) and q-value distribution for SVGs detected by SpatialDE (12,678 genes in total) in the mouse posterior brain dataset.

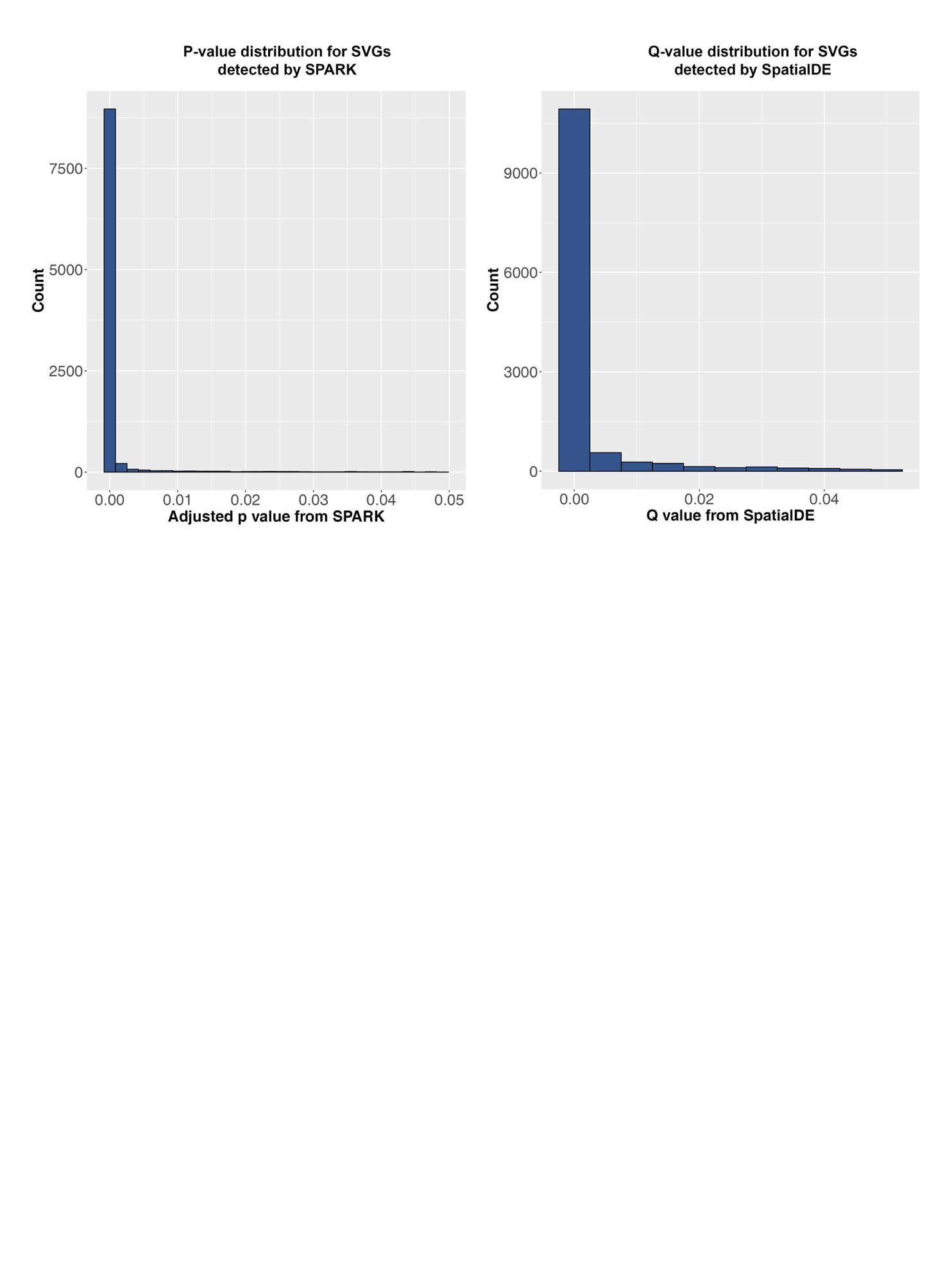

**Supplementary Figure 11.** Gene expression patterns for 40 SVGs randomly selected from the 587 SVGs that were identified by SpaGCN in in the mouse posterior brain dataset.

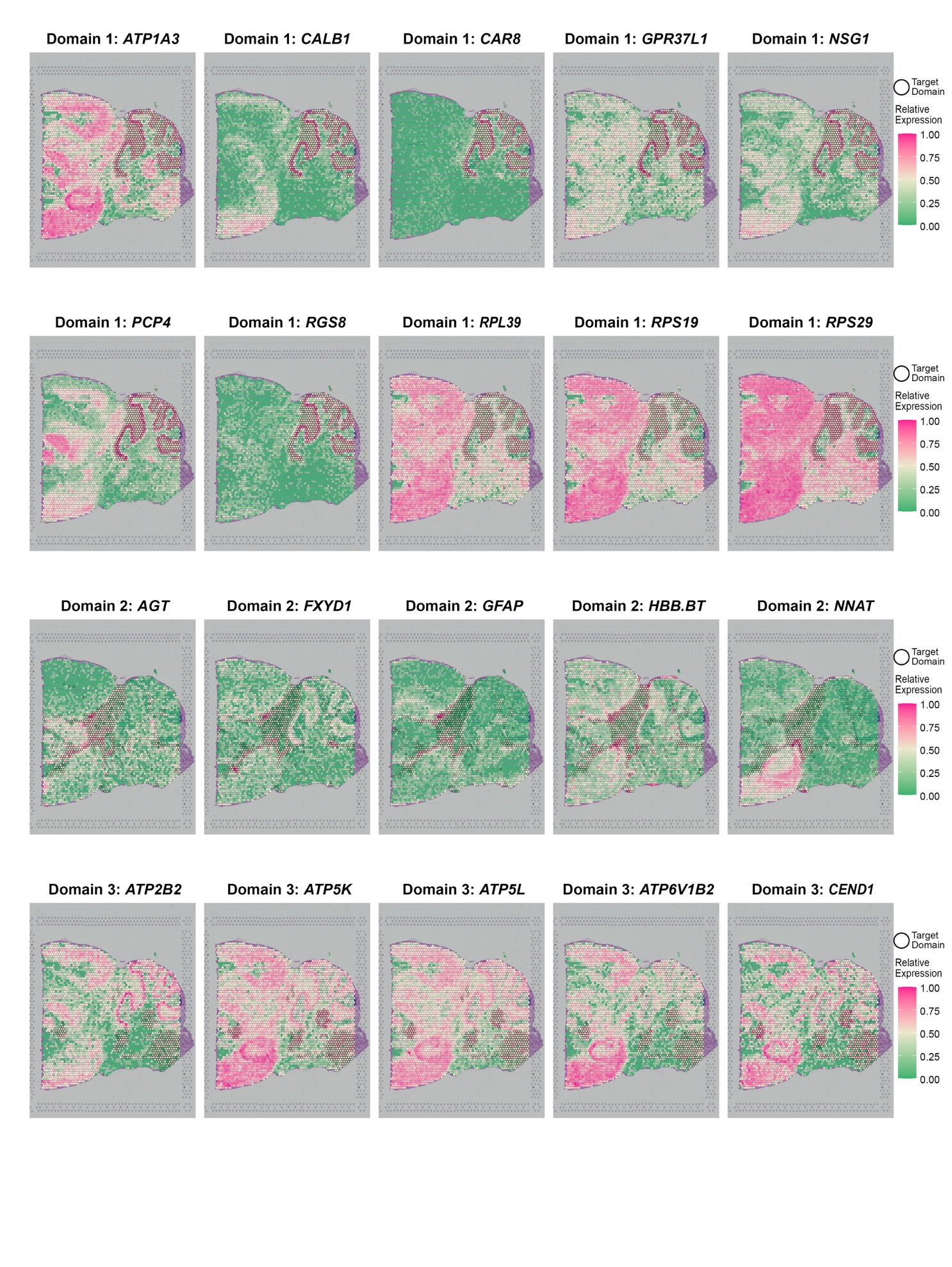

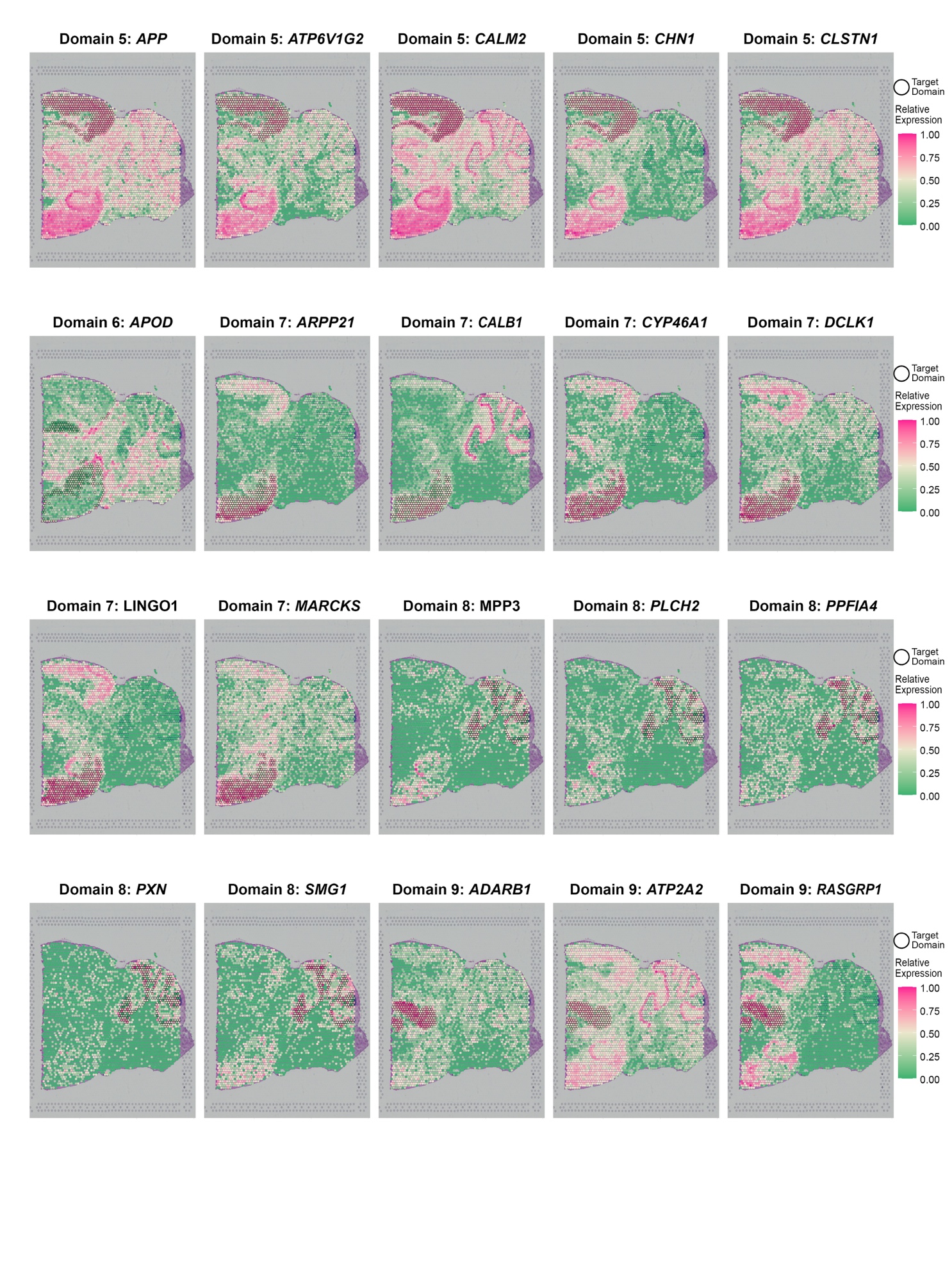

**Supplementary Figure 12.** Transferability of SVGs detected by SapGCN in the mouse posterior brain dataset. SVGs were detected in slice 1 and tested on slice 2.

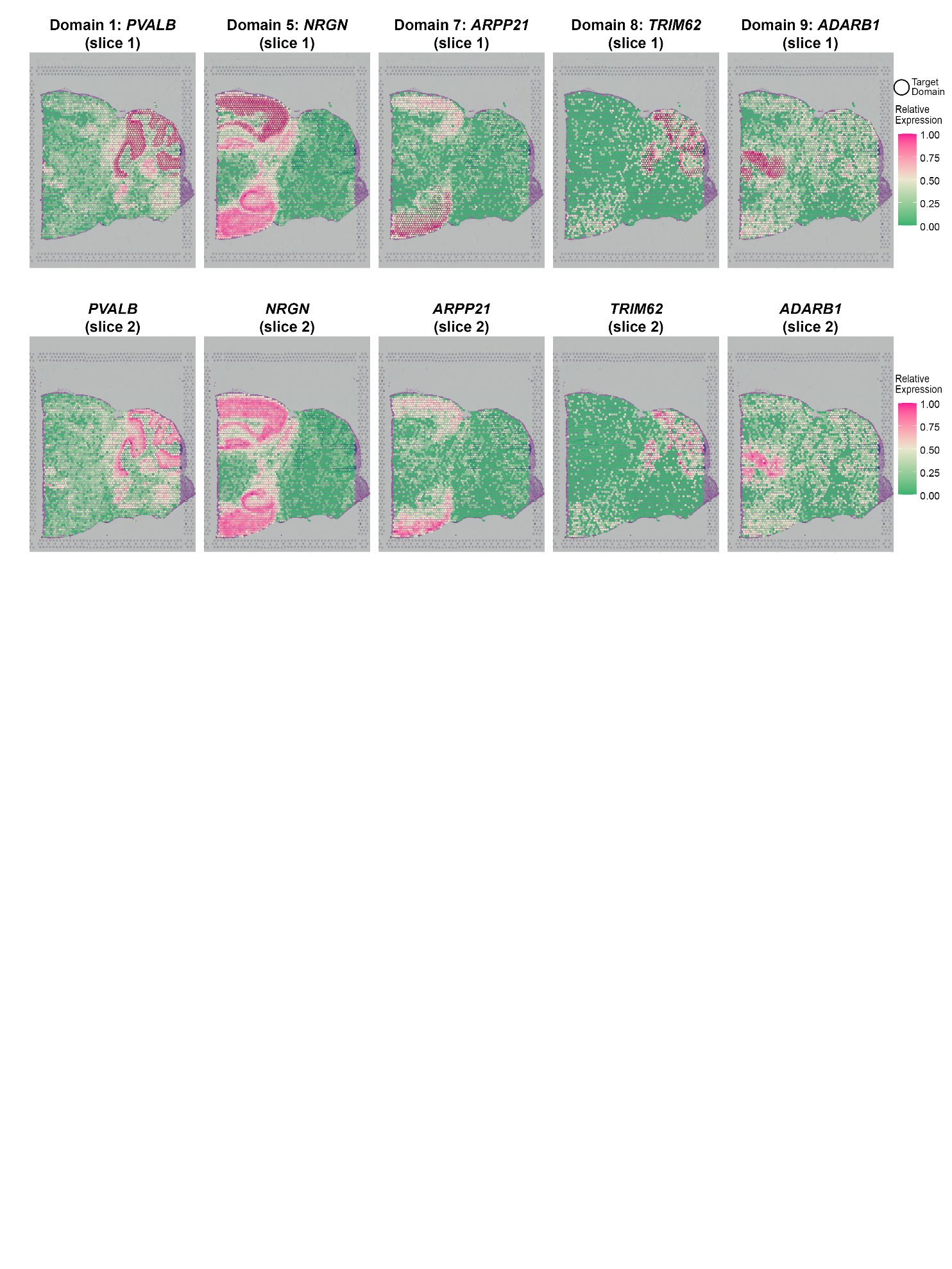

**Supplementary Figure 13.** Transferability of meta gens detected by SapGCN in the mouse posterior brain dataset. Meta genes were detected in slice 1 and tested on slice 2.

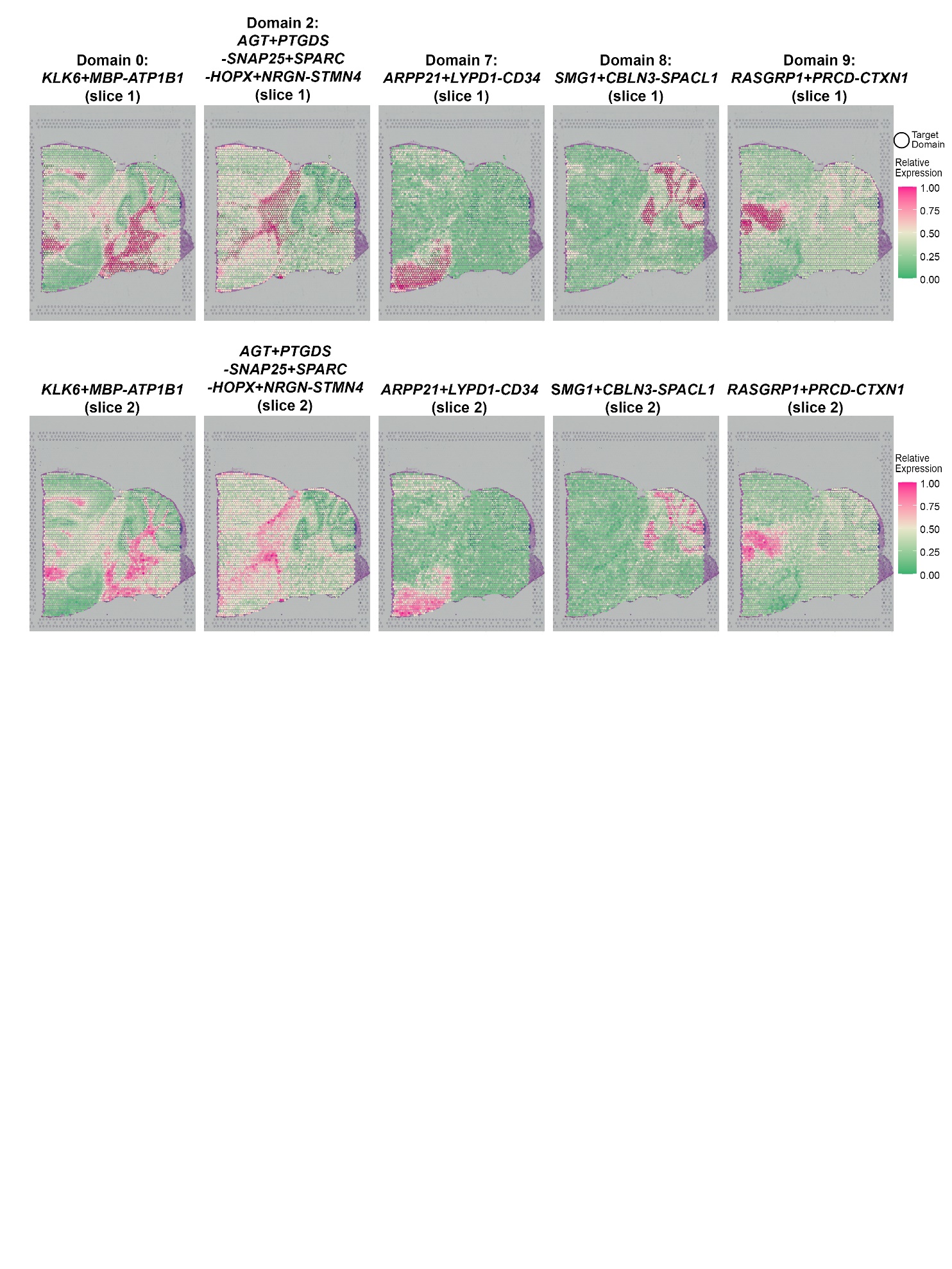

**Supplementary Figure 14.** Gene expression patterns for 20 SVGs randomly selected from the 61 SVGs that were identified by SpaGCN in the LIBD human dorsolateral prefrontal cortex dataset (slice 151673).

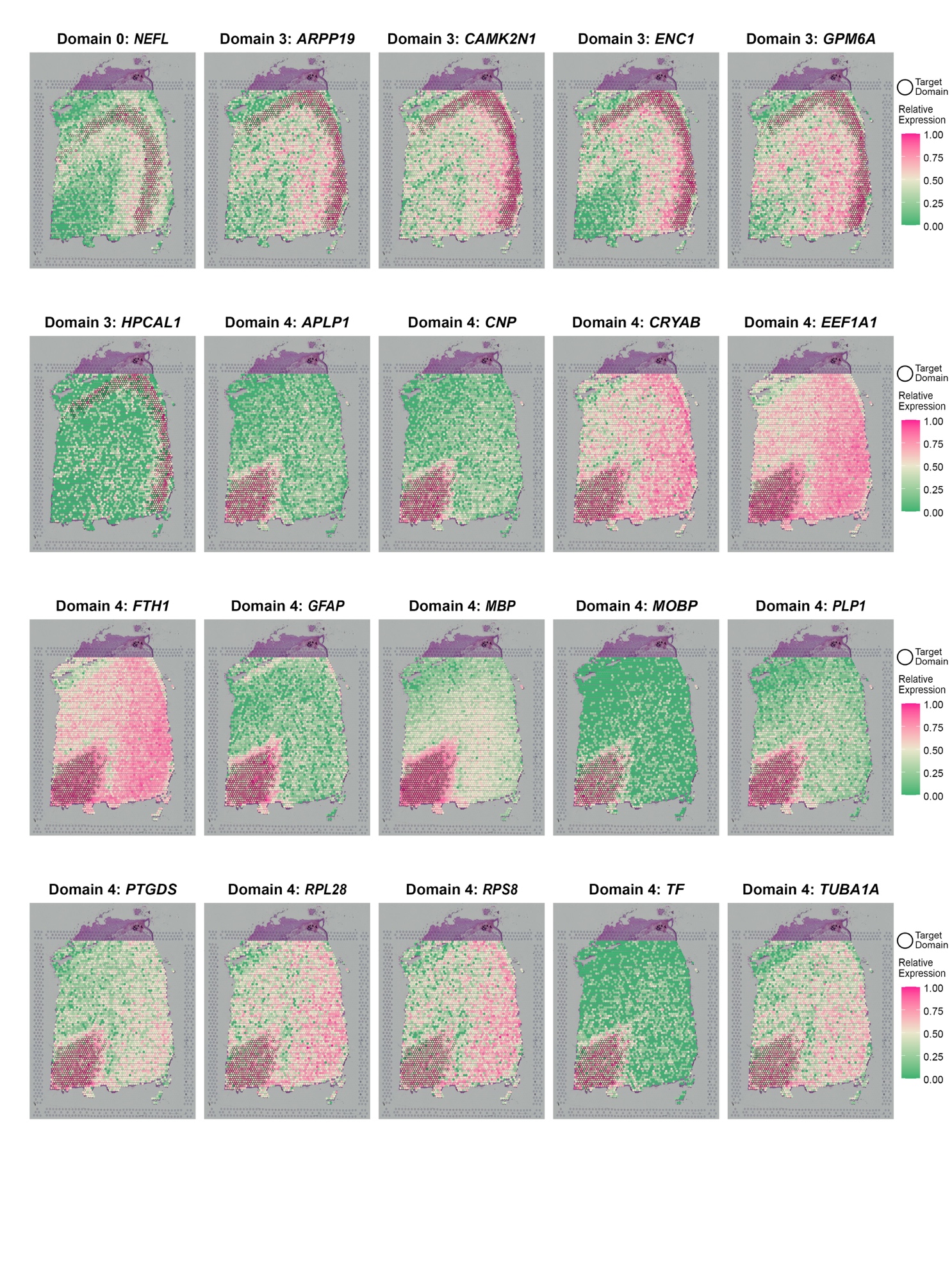

**Supplementary Figure 15.** SVGs detected by SPARK with smaller FDR adjusted p-values do not necessarily show better spatial pattern than genes with larger p-values in the LIBD human dorsolateral prefrontal cortex dataset (slice 151673).

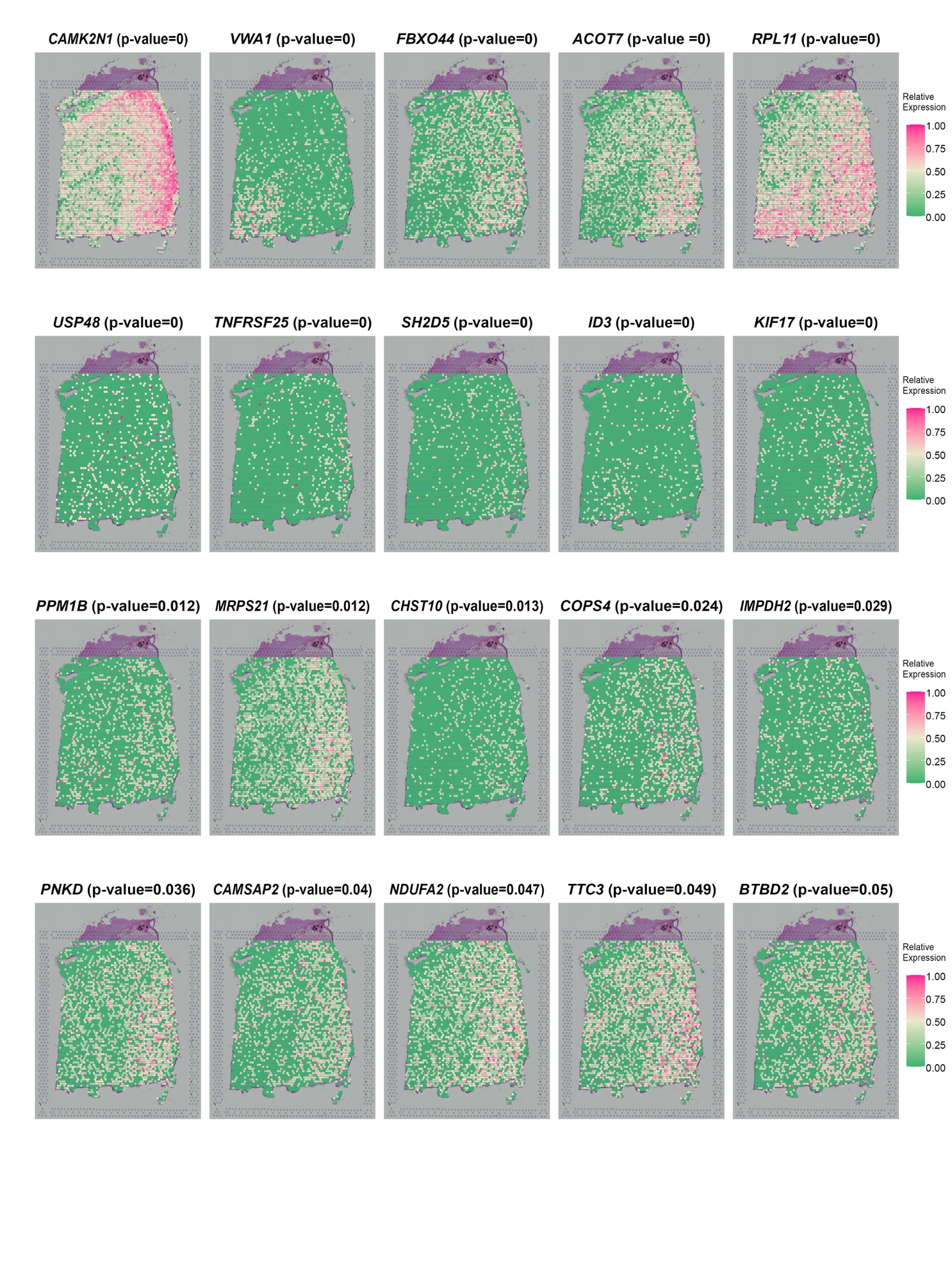

**Supplementary Figure 16.** SVGs detected by SpatialDE with smaller q-values do not necessarily show better spatial pattern than genes with larger q-values in the LIBD human dorsolateral prefrontal cortex data (slice 151673).

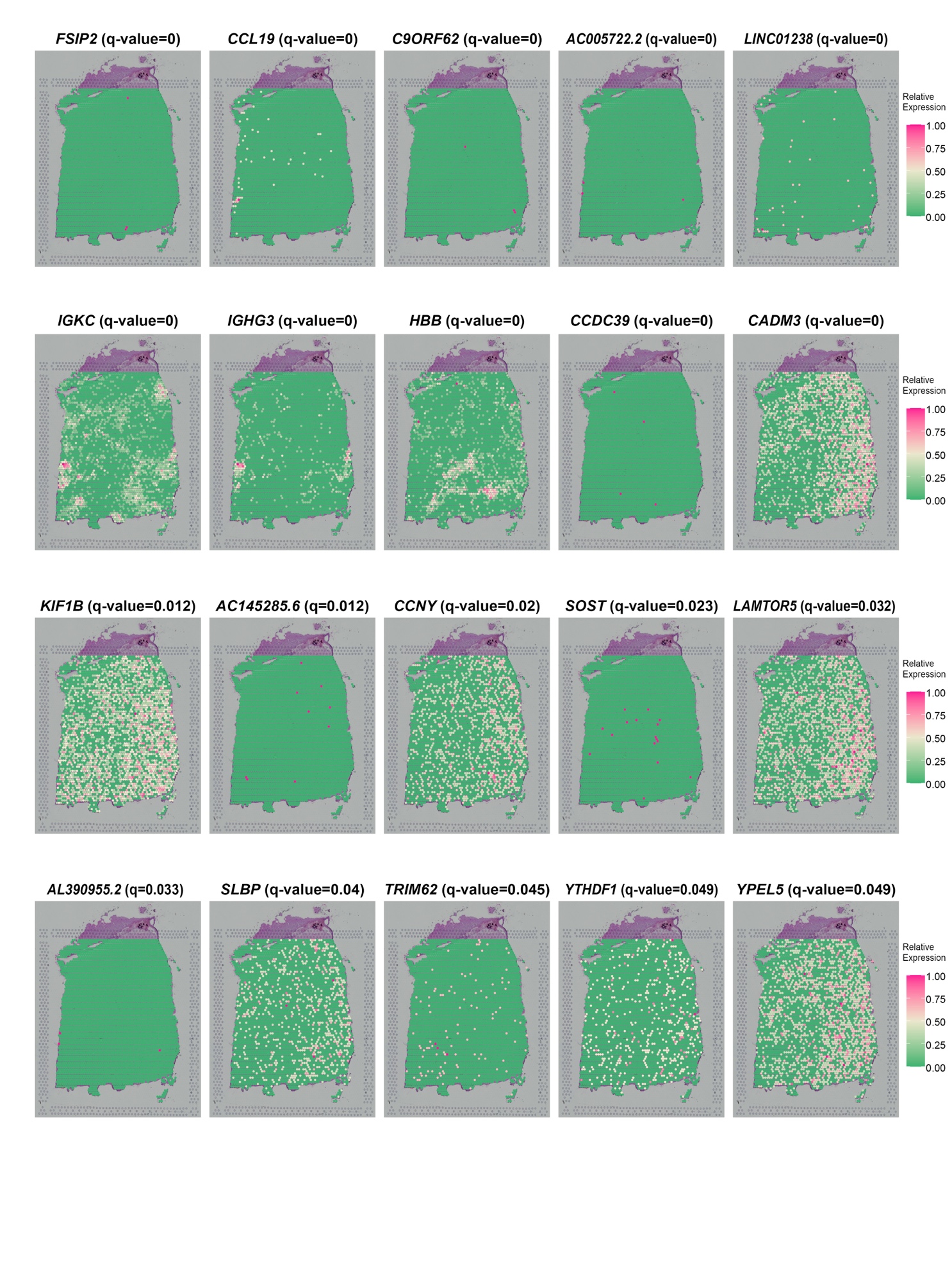

**Supplementary Figure 17.** Venn diagram for SVGs detected by SpaGCN, SpatialDE and SPARK in the LIBD human dorsolateral pre-frontal cortex dataset (slice 151673).

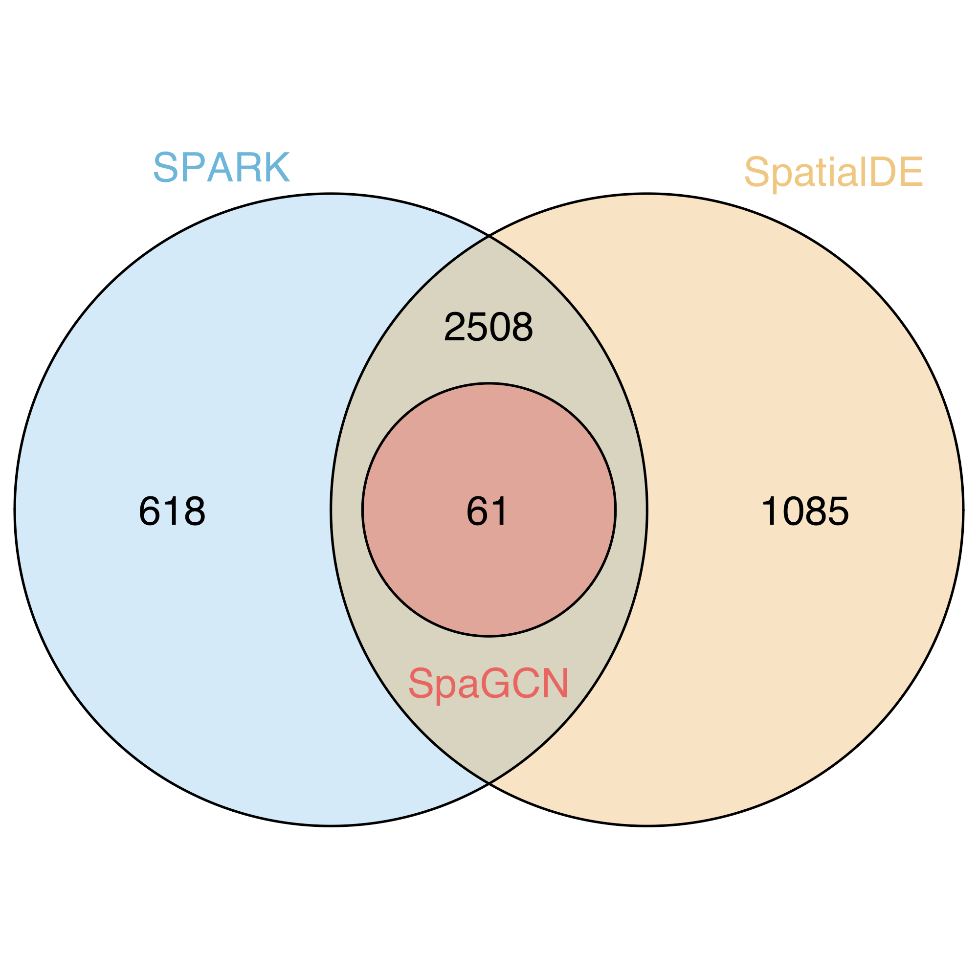

**Supplementary Figure 18.** P-value (FDR adjusted) distribution for SVGs detected by SPARK and q-value distribution for SVGs detected by SpatialDE in the LIBD human dorsolateral pre-frontal cortex dataset (slice 151673).

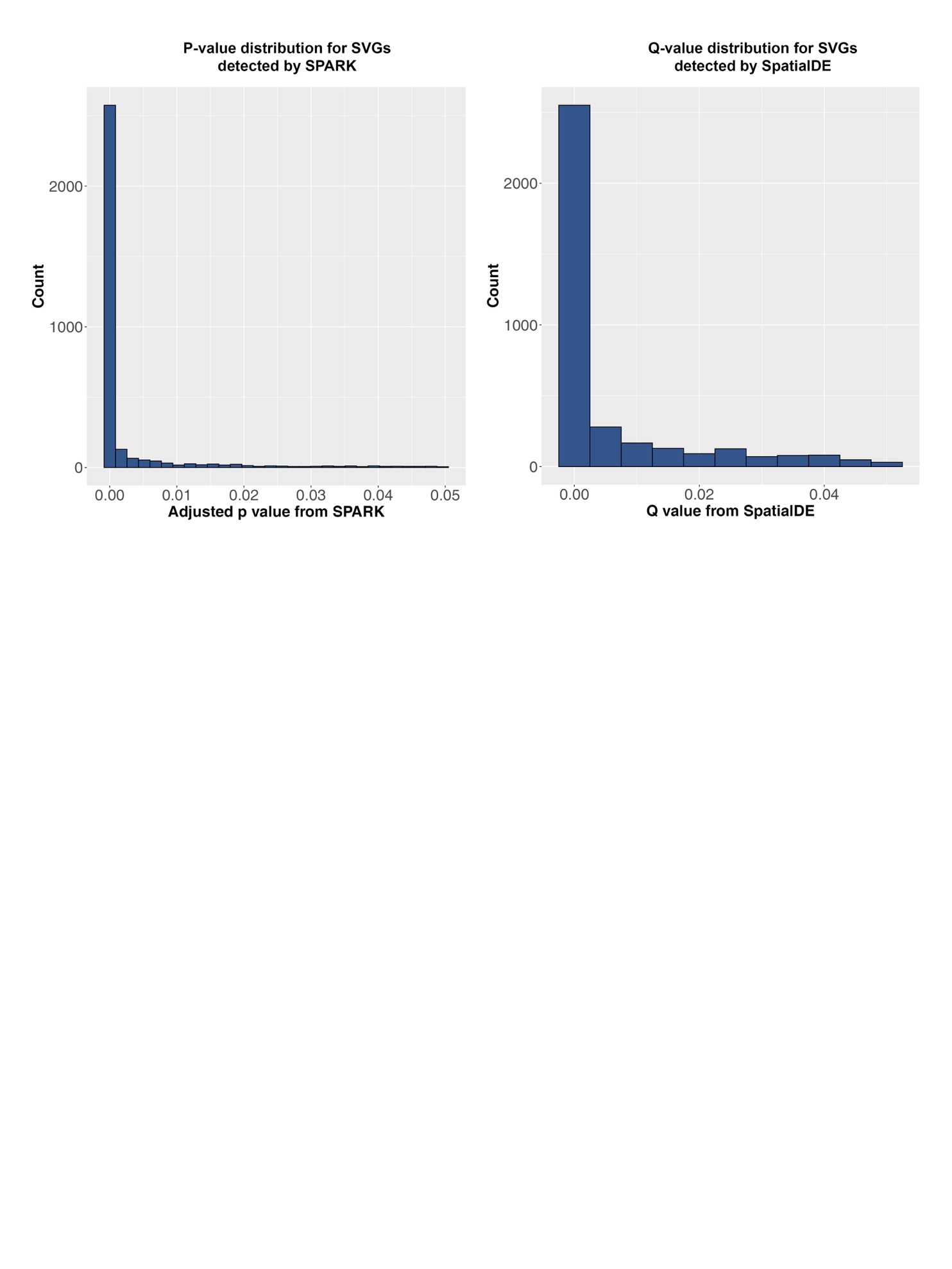

**Supplementary Figure 19.** Transferability of SVGs detected by SapGCN in the LIBD human dorsolateral prefrontal cortex dataset. SVGs were detected in slice 151673 and tested on slice 151676.

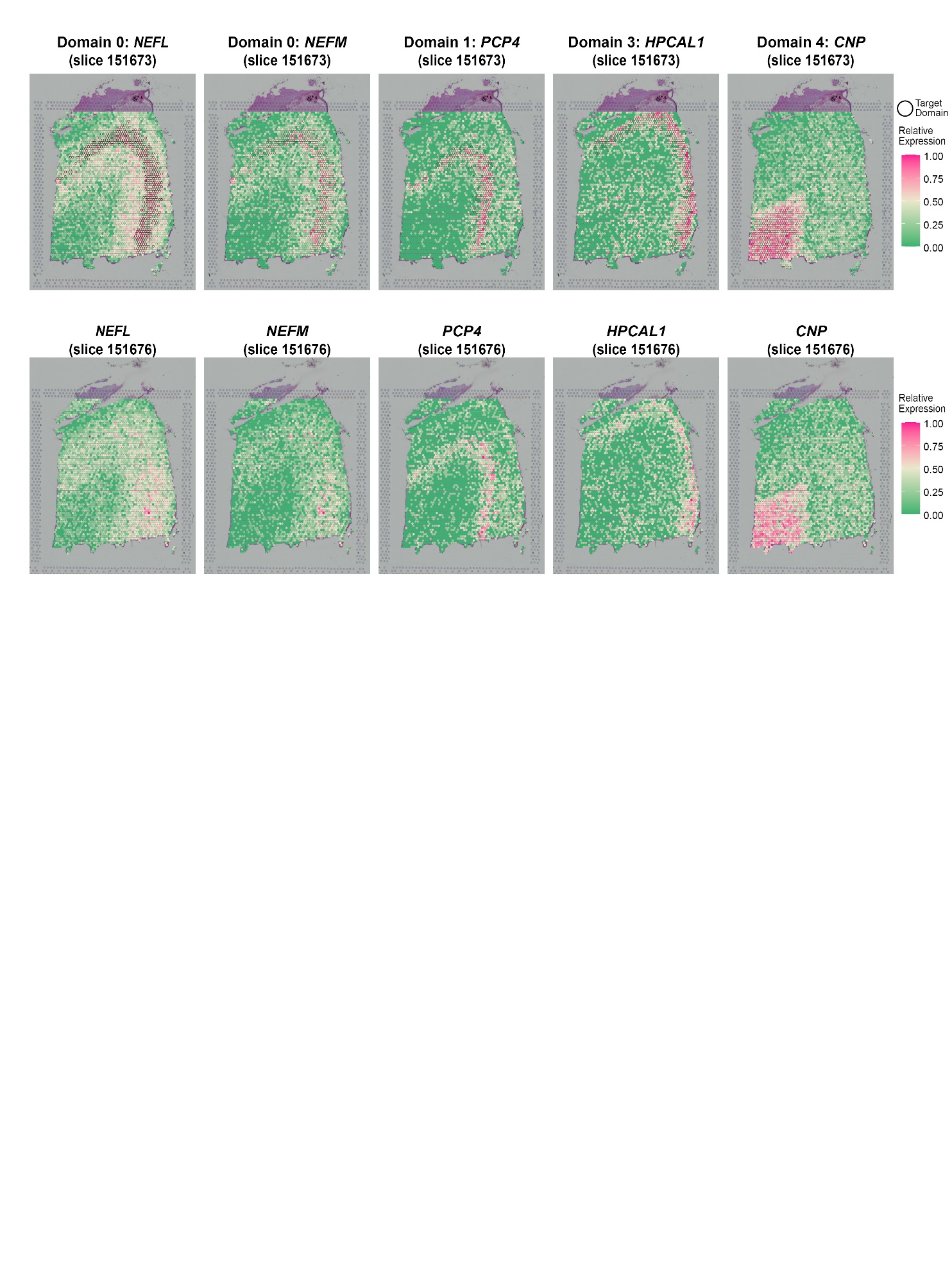

**Supplementary Figure 20.** Transferability of meta gens detected by SapGCN in the LIBD human dorsolateral prefrontal cortex dataset. Meta genes are detected in slice 151673 and tested on slice 151676.

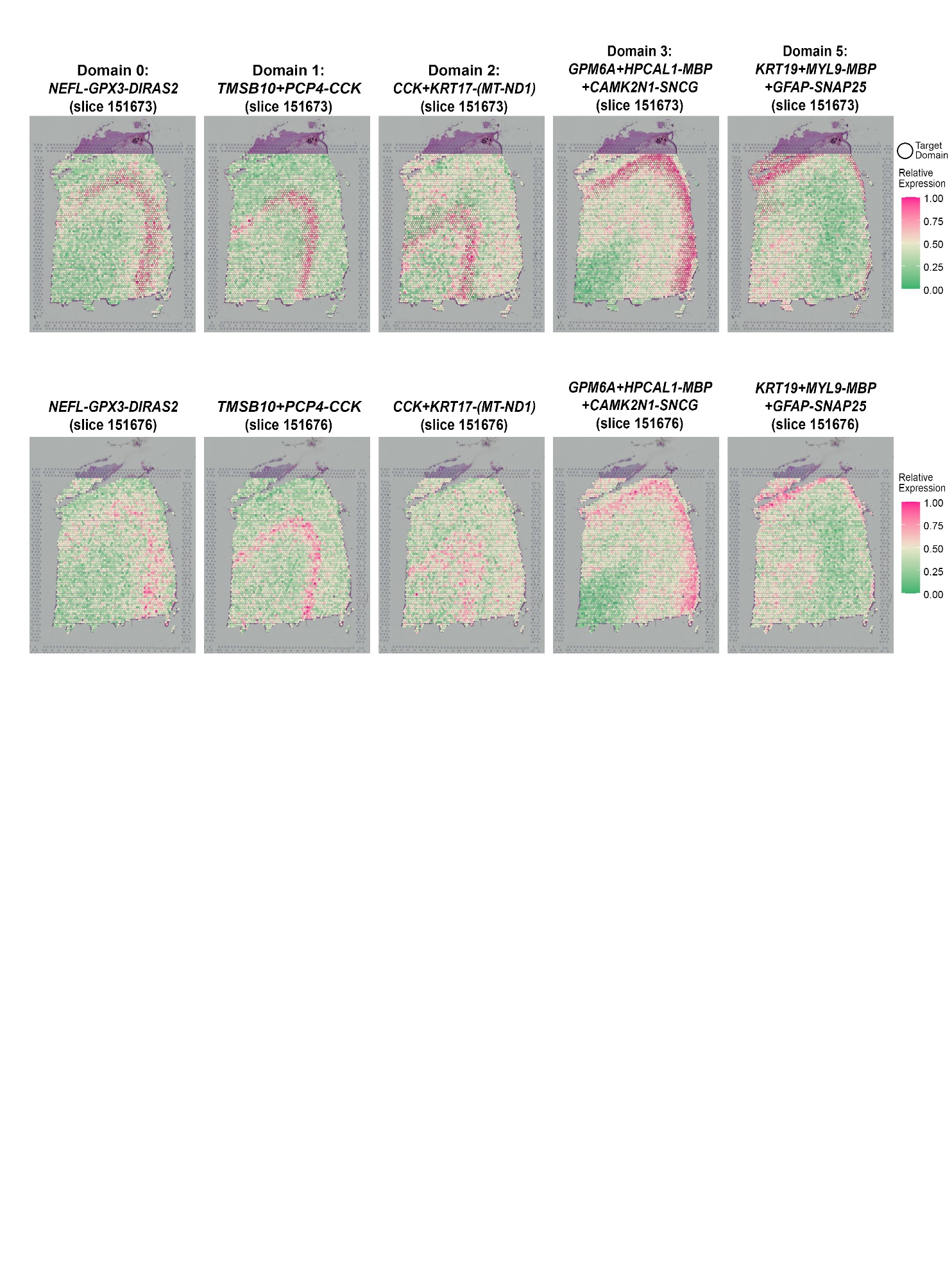

**Supplementary Figure 21.** Gene expression patterns for 10 SVGs that were identified by SpaGCN in the human primary pancreatic cancer tissue.

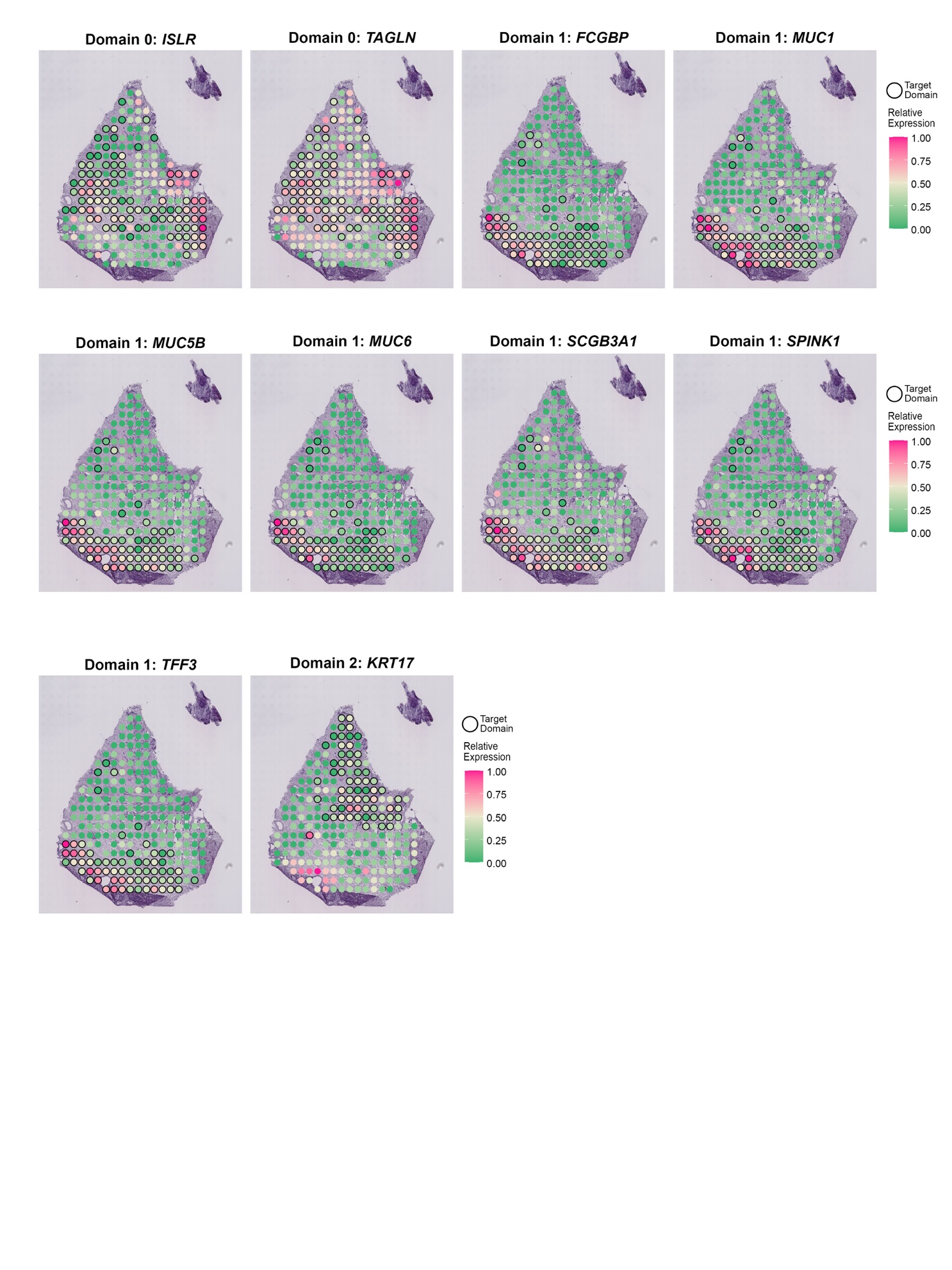

**Supplementary Figure 22.** Venn diagram for SVGs detected SpaGCN, SpatialDE and SPARK in the human primary pancreatic cancer tissue.

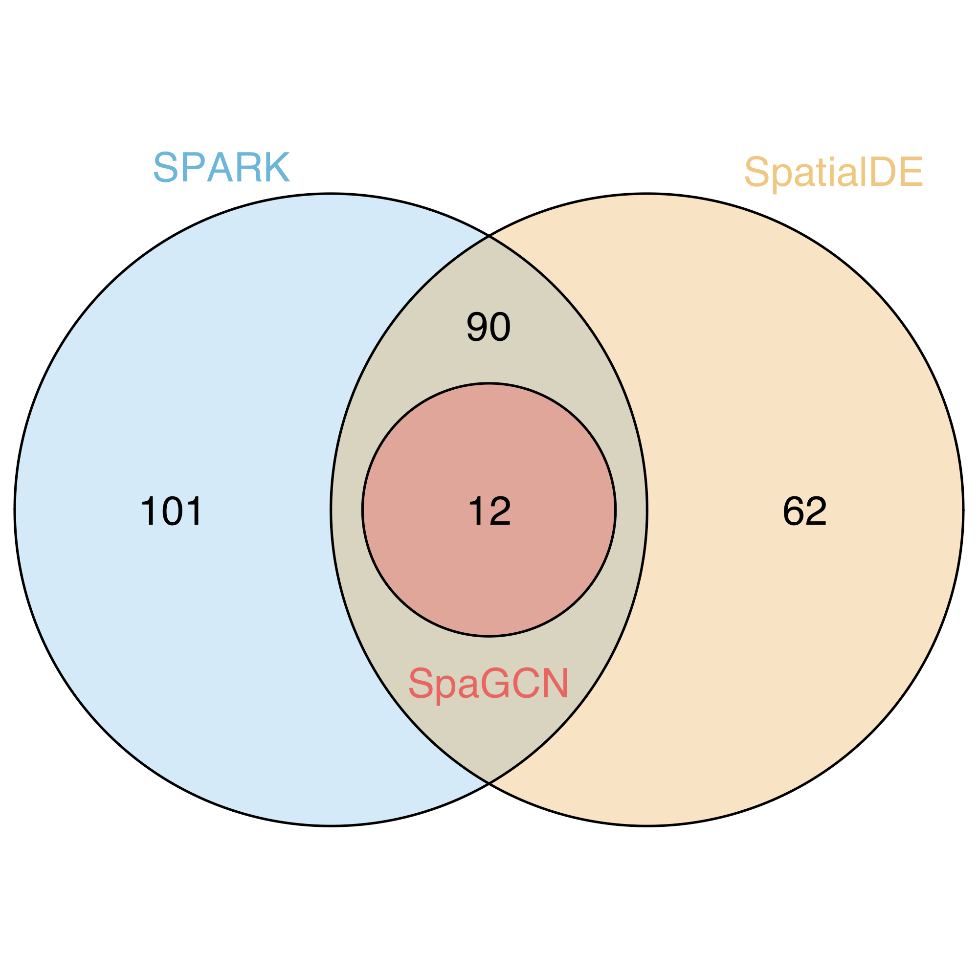

**Supplementary Figure 23.** Gene expression patterns for 16 SVGs that were identified by SpaGCN for domain 2, 3, 7 in the MERFISH mouse hypothalamus data.

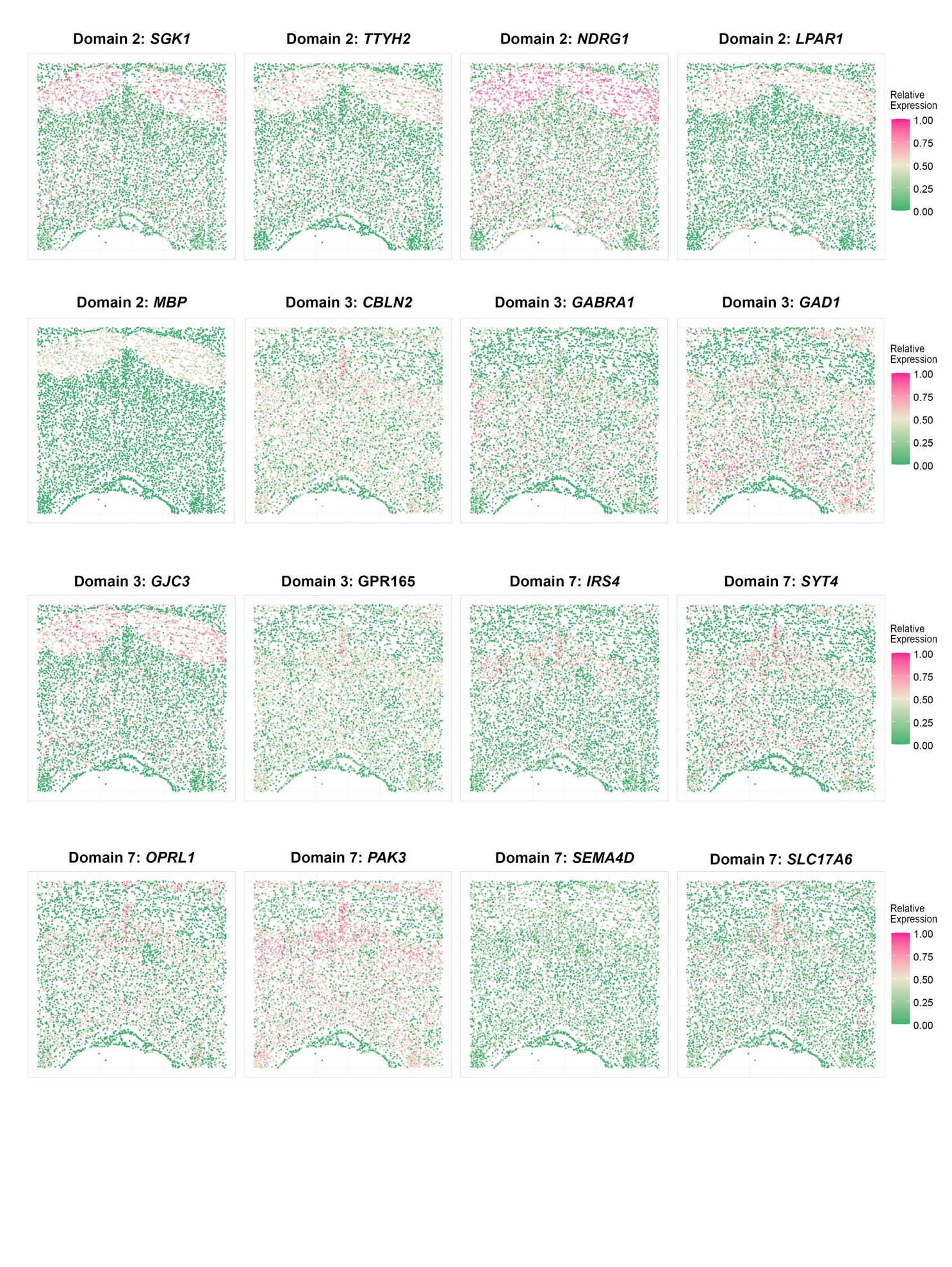

**Supplementary Figure 24.** Computation cost of SpaGCN, SPARK and SpatialDE on the mouse posterior brain dataset.

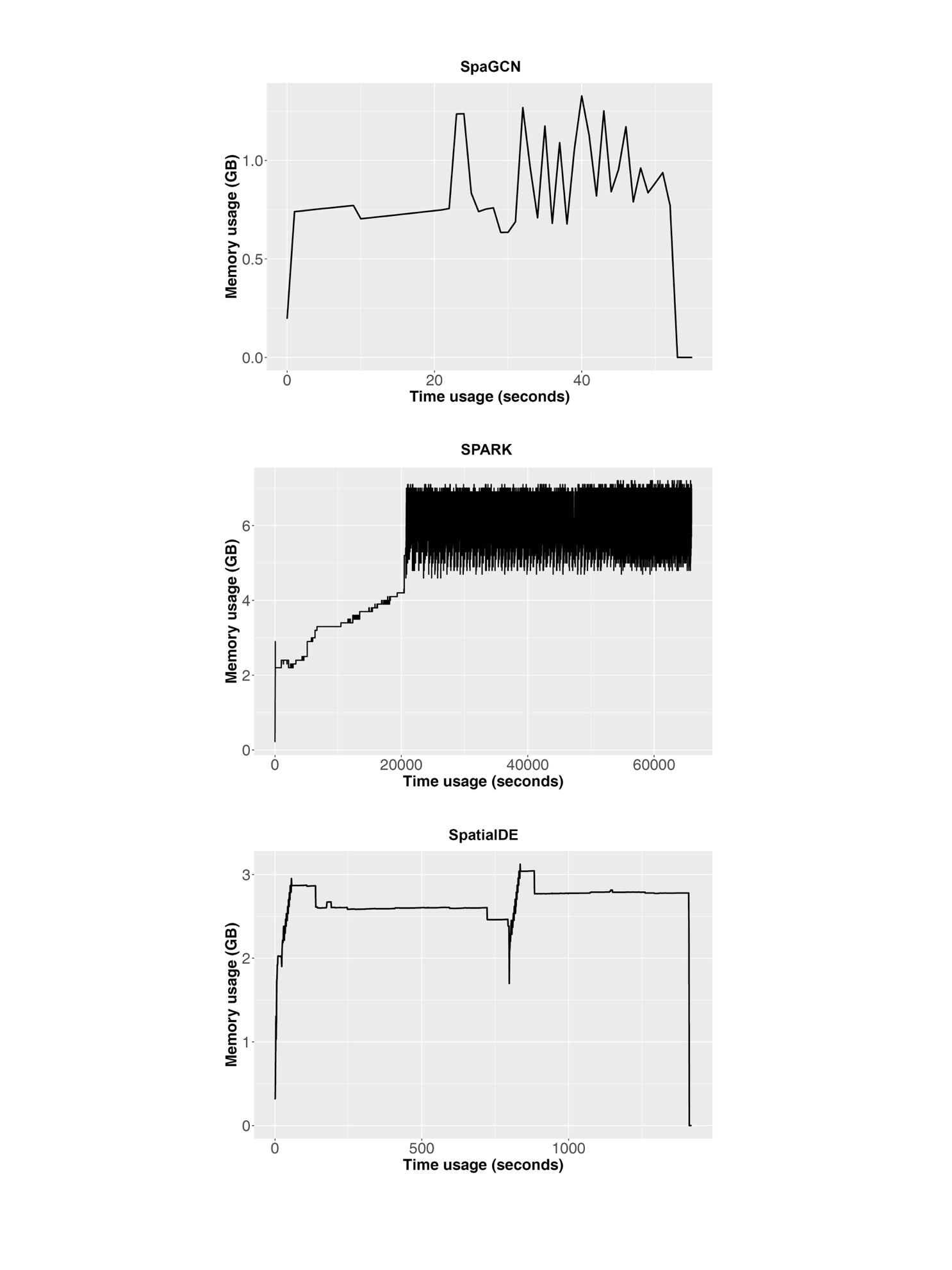

**Supplementary Figure 25.** Illustration of the iterative search of meta gene with spatial gene expression enrichment using the LIBD human dorsolateral pre-frontal cortex dataset (slice 151673). Starting from *KRT8*, SpaGCN creates meta gene 1 to strengthen the target domain specific expression pattern by adding *MYL9* and subtracting *MBP* (meta gene 1 = *KRT8* + *MYL9 – MBP*). This process is repeated to generate meta gene 2 (*KRT8* + *MYL9* – *MBP* + *GFAP* - *SNAP25*) and meta gene 3 (*KRT8* + *MYL9* – *MBP* + *GFAP* – *SNAP25* + *SNAP25* – *PLP1*). SpaGCN stops at meta gene 3 because meta gene 4 (*KRT8* + *MYL9* – *MBP* + *GFAP* – *SNAP25* + *SNAP25* – *PLP1* + *PLP1* – (*MT-CO2*)) does not have a better spatial expression pattern in the target domain.

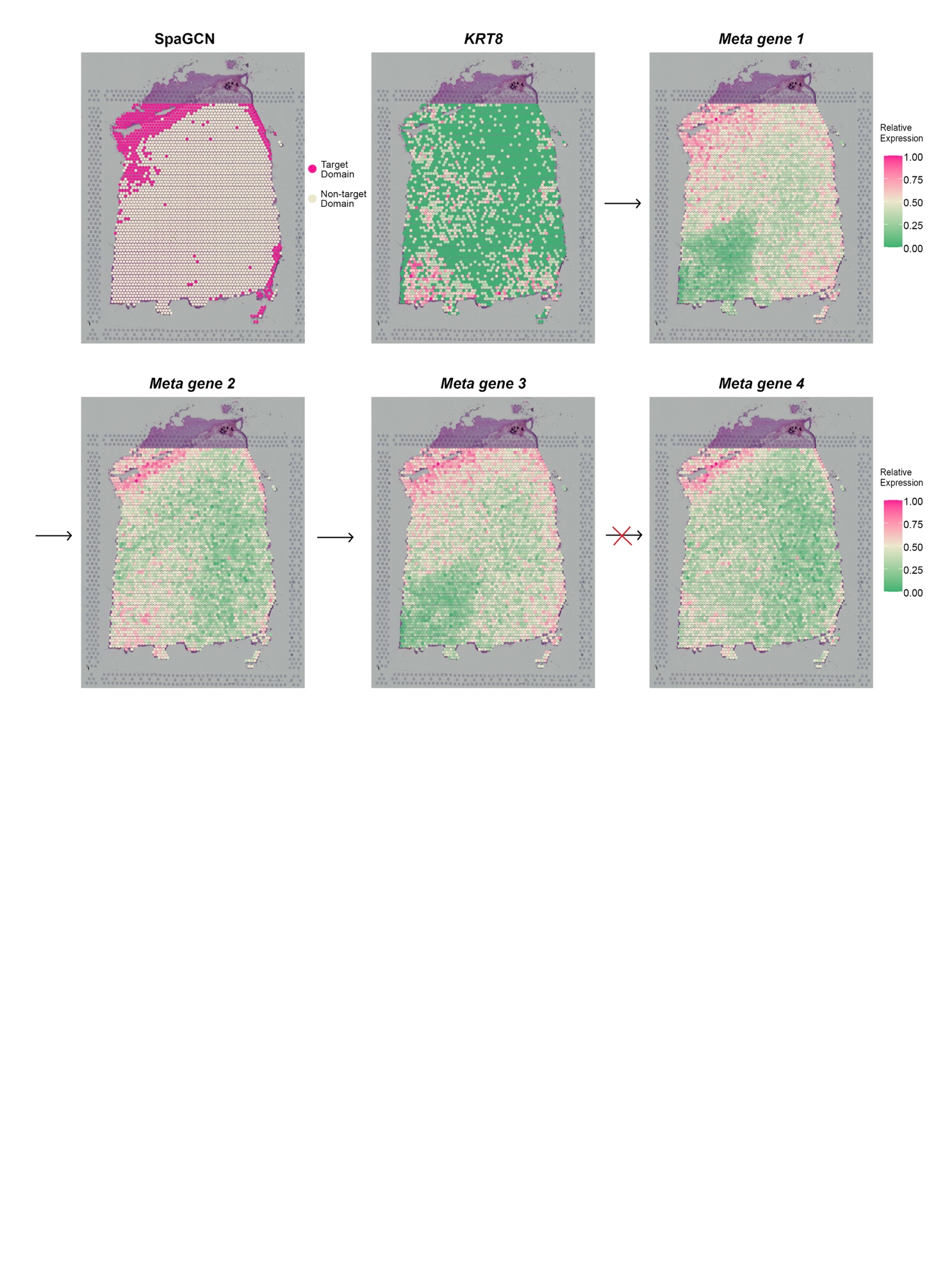
